# Aryl hydrocarbon receptor utilises cellular zinc signals to maintain the gut epithelial barrier

**DOI:** 10.1101/2022.11.03.515052

**Authors:** Xiuchuan Hu, Wenfeng Xiao, Yuxian Lei, Adam Green, Xinyi Lee, Muralidhara Rao Maradana, Yajing Gao, Xueru Xie, Rui Wang, George Chennell, M. Albert Basson, Pete Kille, Wolfgang Maret, Gavin A. Bewick, Yufeng Zhou, Christer Hogstrand

## Abstract

Both zinc and plant-derived ligands of the aryl hydrocarbon receptor (AHR) are dietary components which regulate intestinal epithelial barrier function and protect against Inflammatory Bowel Disease (IBD)^1,2^. Here, we explore whether zinc and AHR pathway are linked using a mouse IBD model with follow-on studies on human and mouse ileum organoids. Our data demonstrate that AHR regulates cellular zinc uptake, and that zinc is an integral part of AHR signalling processes. We show that dietary supplementation in mice with the plant-derived AHR ligand precursor, indole-3-carbinol (I3C), offers a high level of protection against dextran sulfate sodium induced IBD while protection fails in mice with AHR deleted in the intestinal epithelium. AHR agonist treatment is also ineffective in mice with a nutritional zinc deficiency. Experiments in the human Caco-2 cell line and ileum organoids showed that AHR activation increases total cellular zinc and cytosolic free Zn^2+^ concentrations through transcriptional upregulation of several *SLC39* zinc importers. As a consequence, genes for tight junction (TJ) proteins were upregulated in a zinc-dependent manner involving zinc inhibition of signalling to NF-κB and attenuated degradation of TJ proteins through zinc inhibition of calpain activity. Thus, our data indicate that AHR activation by plant-derived dietary ligands improves gut barrier function via zinc-dependent cellular pathways, suggesting that combined dietary supplementation with AHR ligands and zinc might be effective in preventing and treating inflammatory gut disorders.

## Main

The small intestinal epithelium is a single cell-layer composed of many repetitive and self-renewable crypt-villous units^3,4^. Its main functions include allowing the selective passage of nutrients into the body whilst separating the intestinal epithelium from the luminal contents, thereby protecting against pathogenic bacteria, antigens, and other harmful agents^5–8^. The intestinal barrier function is governed by paracellular tight junctions of the epithelium, the mucus layer, and the enteric immune system^9^. The barrier function is compromised in IBD conditions, such as Crohn’s disease and ulcerative colitis, causing leakage and inflammation^7,10^. Conversely, inflammation of the enteric immune system may cause dysregulation of the epithelial barrier and IBD.

Both the aryl hydrocarbon receptor (AHR) and zinc are essential for the development of a fully differentiated intestinal epithelium and maintenance of its integrity and are also involved in regulation of both the innate and adaptive immune responses^11–17^. During zinc deficiency, growth of intestinal stem cells (ISCs) is attenuated, and tight junctions are compromised, resulting in increased epithelial leakiness^18–20^. This may explain why zinc supplementation is widely used to improve intestinal epithelial barrier function and prevent diarrhoea in children as well as in farm animals^13,21^. The AHR is a transcription factor belonging to the Per-ARNT-Sim–p–basic helix-loop-helix protein family^22^. Nutritionally relevant agonists of AHR include indoles mainly found in cruciferous vegetables, such as broccoli and cabbage^23^. Recent research revealed that AHR activation by plant-based dietary ligands in the intestine regulate intestinal stem cell (ISC) proliferation and differentiation, and epithelial barrier function^15,24,25^. Mice with defective AHR signalling in the intestinal epithelium have an increased susceptibility to enteric infection, which could be corrected by dietary supplementation with AHR ligands^26^.

Since both AHR activation and zinc improve intestinal barrier function, we hypothesised that the therapeutic effect of AHR agonists on gut inflammation is dependent on the dietary zinc supply. To test our hypothesis, we used a DSS-induced IBD juvenile mouse model and treated the animals by daily gavage with a plant-derived AHR ligand precursor, indole-3-carbinol (I3C), in combination with zinc-depleted (5 mg/kg), zinc-replete (35 mg/kg), or zinc-supplemented (100 mg/kg) feed as outlined diagrammatically in Fig. 1A. Neither the different Zn diets nor I3C gavage had an effect on body weight without DSS treatment (Extended Data Fig. 1A). DSS treatment caused pronounced body weight loss (Extended Data Fig. 1A), diarrhoea and blood in the faeces (Fig. 1B) with the most severe effects observed in mice fed the zinc-depleted diet. I3C treatment significantly alleviated the body weight loss and intestinal histopathology in DSS-induced IBD for mice on 100 mg/kg or 35 mg/kg zinc diets, but not in the 5 mg/kg zinc diet group. Furthermore, reduced colon length, a marker of intestinal inflammation with very low variability in the DSS-induced IBD model, was less severe in mice treated with I3C in combination with diets containing 35 mg/kg or 100 mg/kg zinc; however, DSS-induced decrease in colon length could not be rescued by I3C in mice fed on the 5 mg/kg zinc diet group (Fig. 1C). DSS-treated mice developed multiple erosive lesions and significant inflammatory responses in colon and ileum, characterized by increased inflammatory cell infiltrations, goblet cell loss, crypt abscess formation and submucosal oedema. I3C-treated mice fed on diets containing 35 or 100 mg/kg zinc displayed substantially lower levels of inflammatory lesions compared with those on the same diets without I3C treatment (Fig. 1D, Extended data Fig. 1B) and this was found to be statistically significant when quantified using blinded scoring (Fig. 1E, Extended data Fig. 1C). However, no benefit of I3C treatment was observed in DSS-exposed mice given the 5 mg/kg zinc diet. Gut barrier integrity is in part maintained by tight junction proteins such as occludin (OCLN) and the mucosal barrier, which is modulated by mucin-2 (MUC2) expression. Expression of *MUC2* has been shown to be reduced in an *in vitro* goblet cell model (HT-29-MTX) following treatment with zinc depleted media^27^ and zinc was found to stimulate secretion of mucus in the fish intestine^28^. Therefore, to evaluate intestinal barrier function, we determined expression levels of OCLN and MUC2 in colon tissues by immunohistochemistry (IHC). The IHC results showed abundant OCLN and MUC2 in the colonic epithelial cells in all conditions without DSS, while they were lower in mice given DSS (Fig 1F, Extended data Fig.1D). I3C treatment of mice exposed to DSS resulted in markedly higher expression of OCLN and MUC2 than mice not treated with I3C, but only in the groups fed on the 35 or 100 mg/kg zinc diets (Fig. 1F). Mice fed on the 5 mg/kg zinc diet did not respond to I3C in terms of maintaining higher OCLN and MUC2 expression following DSS exposure. Together, our results indicate that the therapeutic effect of I3C in DSS-induced IBD model is dependent on sufficient dietary zinc intake.

**Figure 1:**
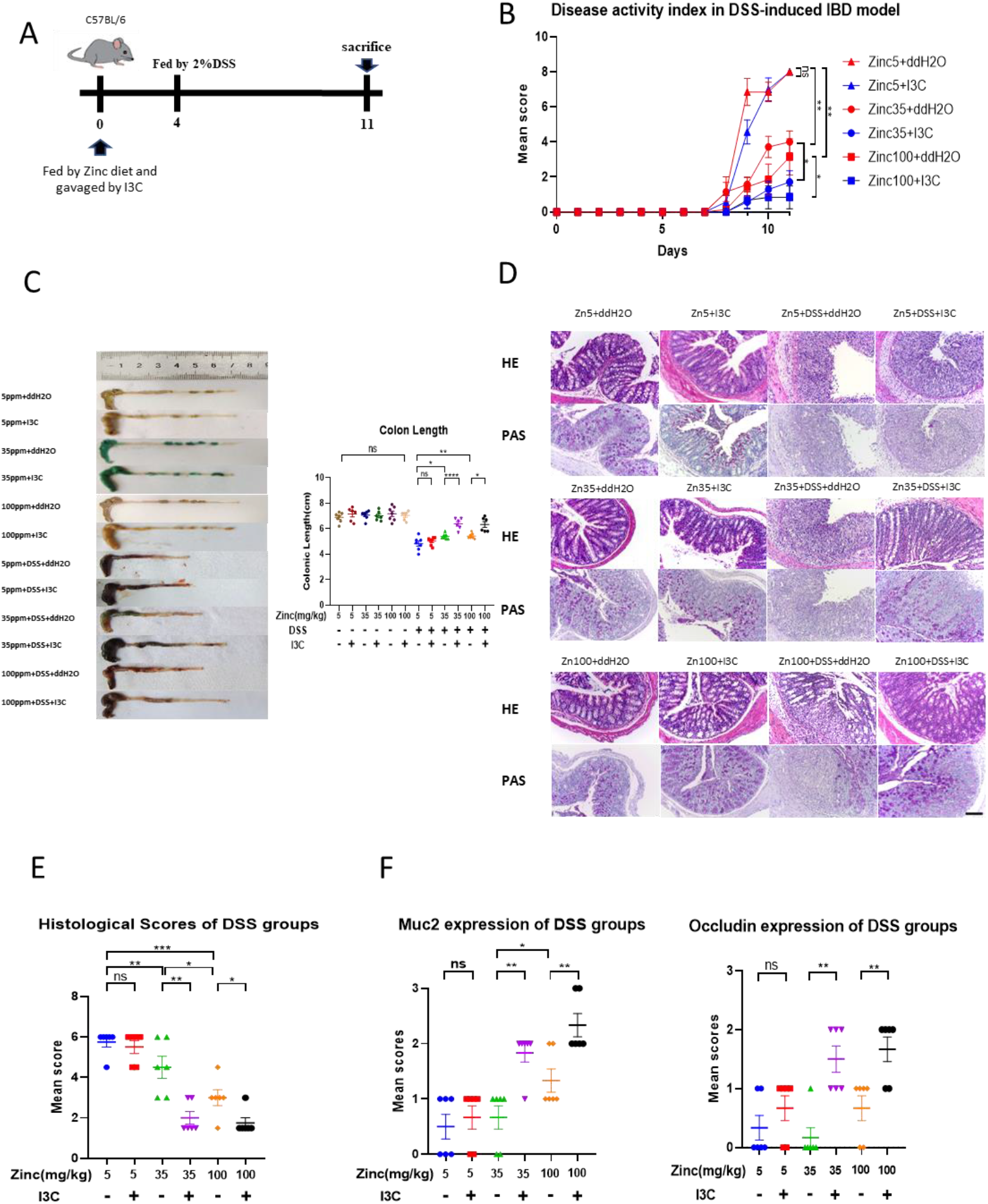
Effects of zinc deficiency and I3C treatment on intestinal integrity in DSS-induced IBD mouse model. (A) Schematic representation of the experimental schedule. Three-week-old C57BL/6J mice were provided with diets containing one of three zinc concentrations (5mg/kg (Zinc5), 35mg/kg (Zinc35) and 100mg/kg (Zinc100) from Days 0-10 with (I3C) or without (ddH2O) I3C given by daily gavage. DSS was administered by the drinking water from Day 4 to Day10. Mice were sacrificed on Day 11. (B) Changes in intestinal disease activity index, based on diarrhoea and bleeding. (C) Colon image (Left) and colon lengths (right) measured on Day 11. (D) Histopathological changes in the colon tissue were examined by H&E and Periodic Acid Schiff^55^ staining (magnification, ×100). Scale bar, 100 μm. (E) Histopathological scores of the colon tissue in DSS treated groups. (F) Immunohistochemistry scoring of MUC2 and Occludin protein expression in colon tissue. Representative data are means ± SEM and n=6~7 in each group. Animal experiments were repeated twice. Statistical analysis was performed using 1-way ANOVA followed by Tukey’s multiple comparison tests. *p<0.05, **p<0.01, ***p<0.001, ****p<0.0001, ns not significant.

To verify if the therapeutic effect of I3C against DSS-induced colitis depended on AHR expression in intestinal epithelial cells, we generated intestinal epithelial-specific AHR knockout mice (villin^cre^ *Ahr*^fl/fl^) and repeated our DSS experiments (Fig. 2A). As predicted, I3C did not mitigate against DSS induced IBD; disease activity (Fig. 2B), weight loss (Extended data Fig. 2A), colon length (Fig. 2C), and histopathology of colon (Fig. 2D, E) and ileum (Extended data Fig. 2B, C) were not different between groups. Interestingly, zinc supplementation alone did not reduce IBD in AHR deficient mice. These results show that intestinal AHR activation by orally administered I3C substantially ameliorates IBD in the DSS mouse model, but the efficacy of the treatment is dependent on a zinc-sufficient diet and is most effective together with zinc supplementation.

**Figure 2:**
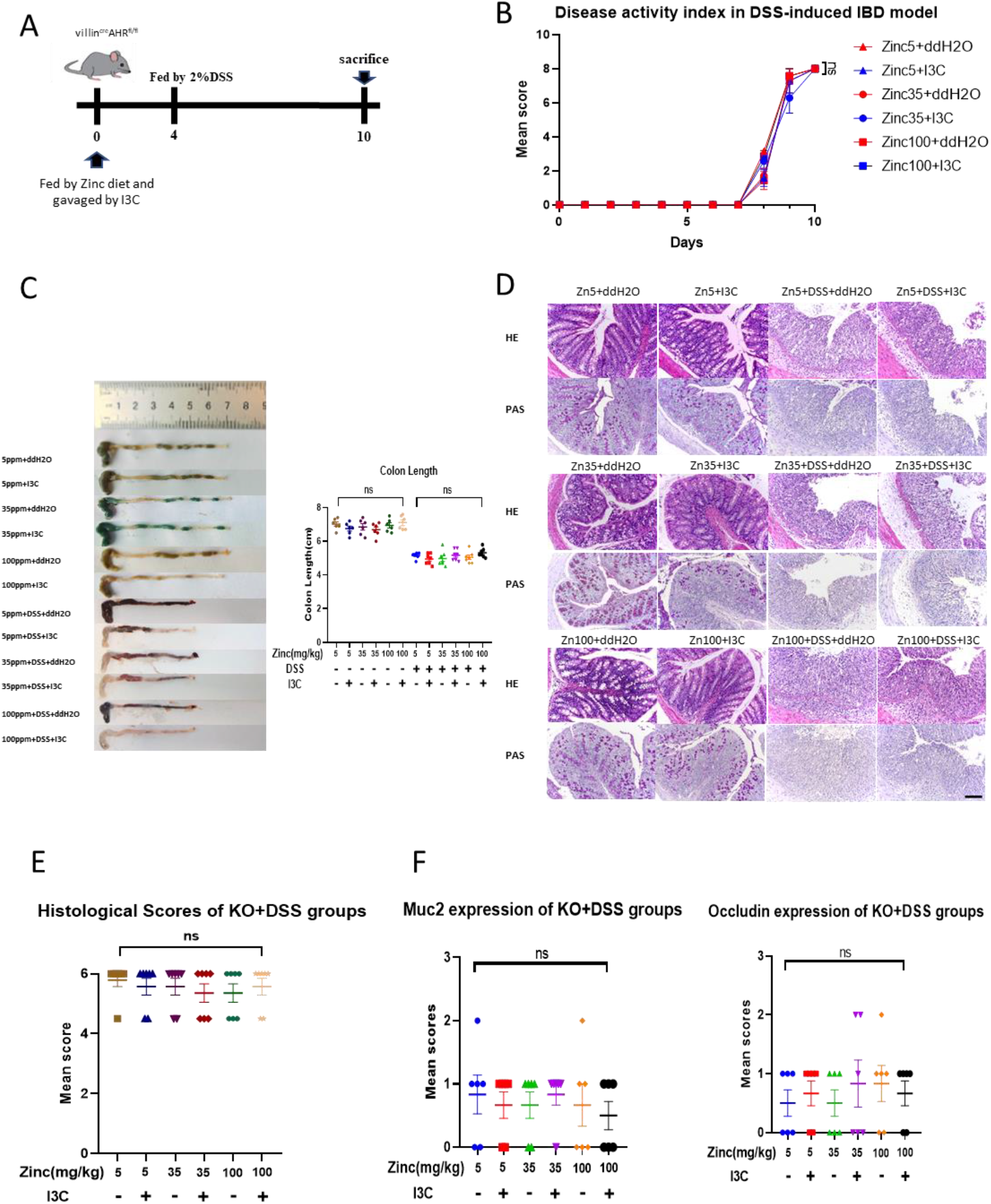
Effects of zinc deficiency and I3C treatment on intestinal integrity in DSS-induced IBD model in Vil1-Ahr KO mice. (A) Schematic representation of the experimental schedule. Three-week-old villin^cre^ *Ahr*^fl/fl^ mice on C57BL/6J background were provided with diets containing one of three zinc concentrations (5 mg/kg (Zinc5), 35 mg/kg (Zinc35) and 100 mg/kg (Zinc100) from Days 0-10 with (I3C) or without (ddH2O) I3C given by daily gavage. DSS was administered by the drinking water from Day 4 to Day10. Mice were sacrificed on Day 11. (B) Changes in intestinal disease activity index, based on diarrhoea and bleeding. (C) Colon image (Left) and colon lengths (right) measured on Day 11. (D) Histopathological changes in the colon tissue were examined by H&E and Periodic Acid Schiff^55^ staining (magnification, ×100). Scale bar, 100 μm. (E) Histopathological scores of the colon tissue in DSS treated groups. (F) Immunohistochemistry scoring of MUC2 and Occludin protein expression in colon tissue. Representative data are means ± SEM and n=6~7 in each group. Animal experiments were repeated twice. Statistical analysis was performed using 1-way ANOVA followed by Tukey’s multiple comparison tests. *p<0.05, **p<0.01, ***p<0.001, ****p<0.0001, ns not significant.

We considered that some of the effects of I3C and zinc treatment observed might have been mediated by influence of these treatments on gut microbiota. We therefore performed 16s rRNA sequencing of the intestinal content of four mice from each DSS group. Dietary zinc concentrations and I3C administration did not affect alpha or beta diversity of species in WT mice, but there was a small increase in alpha diversity of microbes with increasing dietary zinc levels in DSS treated villin^cre^*Ahr*^fl/fl^ mice (Fig. 3A, B). There was also a higher alpha diversity in villin^cre^*Ahr*^fl/fl^ mice compared with the WT, independent of zinc concentration in the diet and this could be attributed to an increase in abundance of several pathogenic species (Fig. 3C-E). We conclude from this analysis that changes in microbiota cannot explain the beneficial effect of zinc and I3C co-treatment on mitigation of DSS-induced pathology.

**Figure 3:**
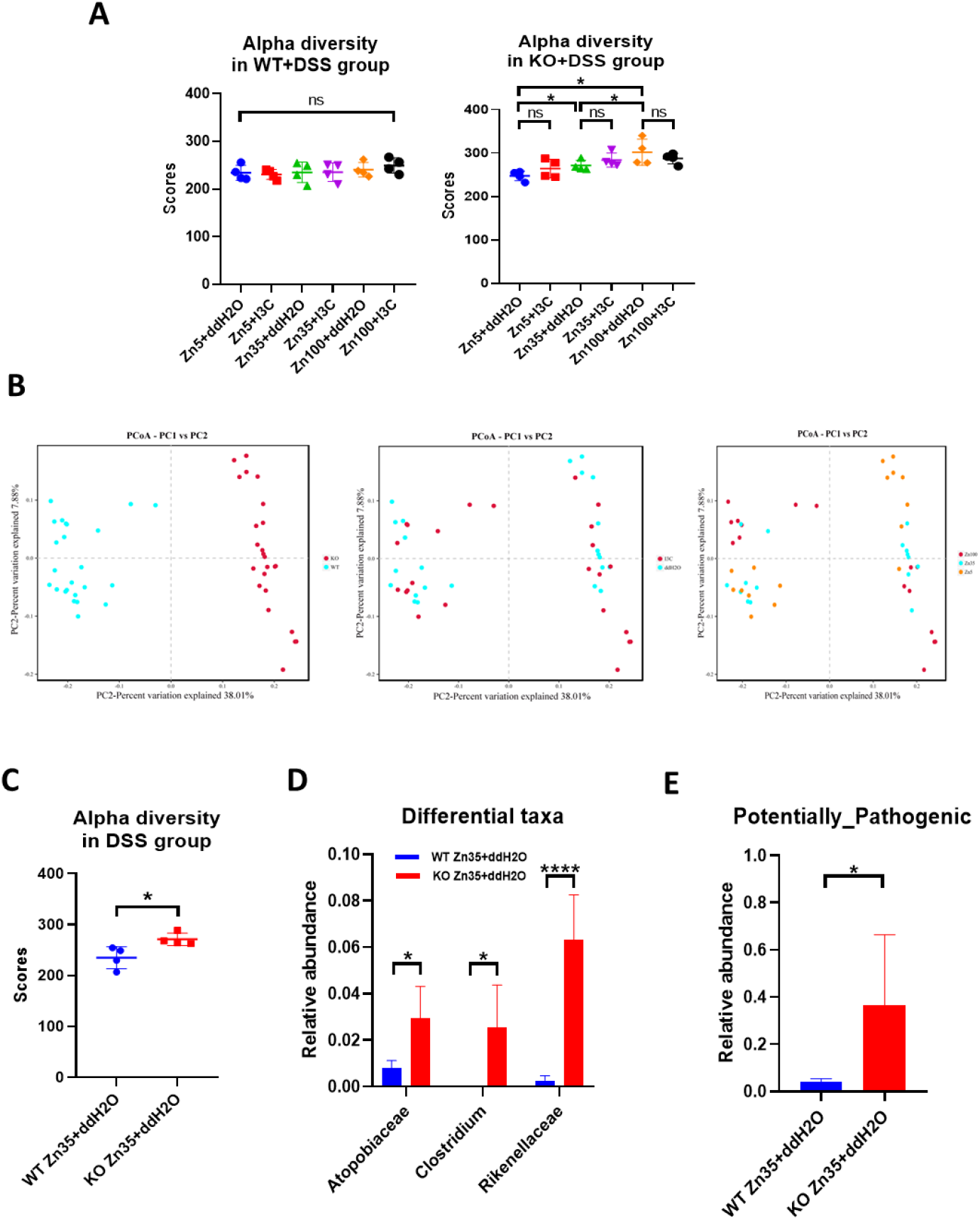
Analysis of intestinal microbiome in DSS-treated WT and Vil1-AhR KO mice. Three-week-old C57BL/6J (WT) mice and villin^cre^ *Ahr*^fl/fl^ mice (KO) on C57BL/6J background were provided with diets with one of three zinc concentrations (5 mg/kg (Zinc5), 35 mg/kg (Zinc35) and 100 mg/kg (Zinc100) from Days 0-10 with (I3C) or without (ddH2O) I3C given by daily gavage. DSS was administered by the drinking water from Day 4 to Day10. Mice were sacrificed on Day 11 and intestinal contents sampled for microbiome profiling. (A) Alpha diversity analysis of DSS-treated WT and vilhn^cre^Ahr^fl/fl^ groups. (B) Beta diversity analysis of WT and villin^cre^Ahr^fl/fl^ groups. Each plot is coloured according to the variables (WT or KO, I3C, Zn). PC1, PC2 represent the top two principal components that captures most of the differences in diversity. (C) Alpha diversity analysis between DSS-treated WT and villin^cre^AHR^fl/fl^ groups, both fed on the diet with intermediate zinc concentration (35 mg/kg (Zn35)) and not administered I3C (ddH2O). (D) The three most differing prokaryotic intestinal taxa between between DSS-treated WT and villin^cre^AHR^fl/fl^ groups, both fed on the diet with intermediate zinc concentration (35 mg/kg (Zn35)) and not administered I3C (ddH2O). (E) Potential pathogenicity prediction of DSS-treated WT and villin^cre^AHR^fl/fl^ groups, both fed on the diet with intermediate zinc concentration (35 mg/kg (Zn35)) and not administered I3C (ddH2O). Representative data are means ± SEM and n=4 in each group. Statistical analysis of the data was performed using 1-way ANOVA followed by Tukey’s multiple comparison tests (Fig. A) and unpaired t-tests (Fig. C, D, E). *p<0.05, **p<0.01, ***p<0.001, ****p<0.0001, ns not significant.

Next, we used *in vitro* models to explore the relevance of our findings in human. We used differentiated Caco-2 cells grown into epithelia which differentiated into enterocytes with characteristics similar to those of small intestine enterocytes^29^, and human ileum organoids. Since I3C is a pro-ligand of AHR and does not function without conversion in the gut, we used FICZ as the AHR agonist in our *in vitro* experiments. To test if AHR activation requires zinc to promote epithelial resistance and reduce paracellular permeability we grew Caco-2 cells on inserts in a Transwell^®^ system. With this system we could mimic the barrier function of the epithelium of the human small intestine and measured its trans-epithelial electrical resistance (TEER) and permeability (leakage of FITC-Dextran 4000) employing different experimental conditions to stimulate a healthy gut, inflammation (DSS treatment), and hypoxia. Addition of zinc or FICZ alone increased electrical resistance and reduced permeability in all experimental conditions but the effect was greater when both agents were added together (Fig. 4A; Extended data Fig. 3A-E). The beneficial effect of the combined treatment with zinc and FICZ on epithelial tightness was particularly strong in the two models of compromised epithelia through exposure to DSS (Fig. 4A, Extended data Fig. 3D) or hypoxia (Extended data Fig. 3B, E) although it was also significant in unchallenged cells representing a healthy intestinal epithelium (Extended data Fig. 3A, C). Human ileum organoids were used to investigate the therapeutic properties of FICZ and zinc on intestinal epithelium barrier function in a system with closer resemblance to the human intestine. Also in this organoid system, the combined treatment of zinc and FICZ reduced epithelial permeability in an additive way (Fig. 4B, Extended data Fig. 3F). Thus, treatment with zinc and FICZ in combination markedly improves barrier function of human Caco-2 cells and ileum organoids.

**Figure 4:**
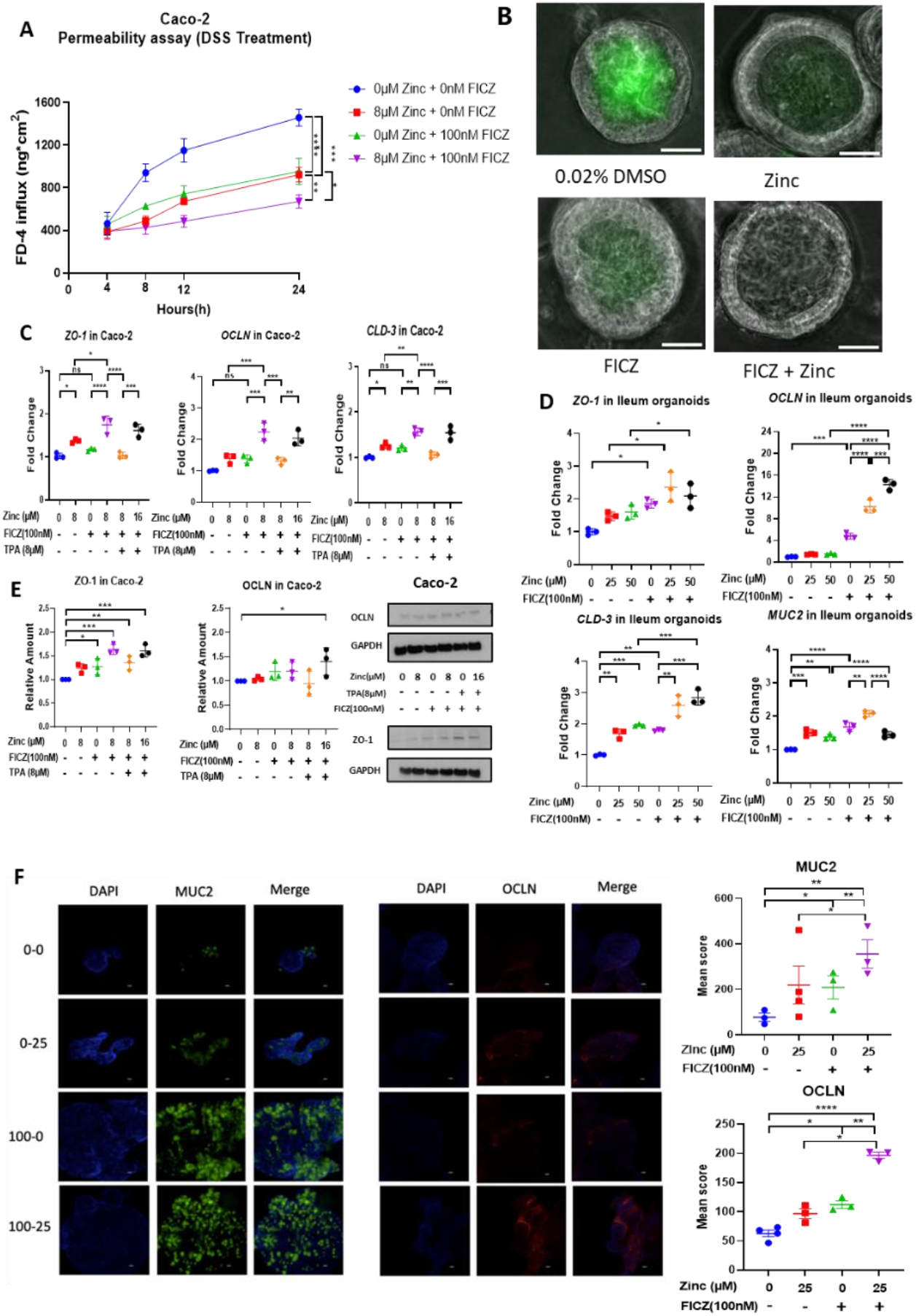
Combined treatment of AHR agonist FICZ and zinc promotes epithelial barrier function in Caco-2 cells and human ileum organoids. (A) Caco-2 cells were grown to an epithelium in a Transwell^®^ system. At the start of the experiment, the medium in the apical compartment was replaced with MEM containing FITC-dextran 4000 (Merck) together with 3% DSS, 0 or 8 μM zinc, and 0 or 100 nM FICZ as indicated in the figure. Permeability was measured by sampling the medium in the basal compartment and measurement of FITC fluorescence over 24 hours. (B) Human ileum organoids were challenged with 60 μM EDTA in presence of FITC-Dextran in a medium containing zinc and/or FICZ. Leakage into the lumen was assessed after 24 h through appearance of FITC fluorescence. (C) Expression of mRNA for tight junction proteins as measured by qPCR in Caco-2 cells treated with zinc/FICZ/TPA for 24 h as indicated in the figure. (D) Expression of mRNA for tight junction proteins and MUC2 as measured by qPCR in human ileum organoids treated with zinc and/or FICZ for 24 h as indicated in the figure. (E) Abundance of tight junction proteins, as measured by western blot, in Caco-2 cells treated with Zinc/FICZ/TPA for 24 h as indicated in the figure. (F) Immunocytochemistry (left) showing MUC2 and OCLN protein levels in human ileum organoids treated with 0 or 100 nM FICZ and/or 0 or 25 μM zinc for 24 h, as indicated in the figure. Fluorescence levels were quantified relative to DAPI using Fiji and numeric data are shown (right). Statistical analysis of the data was performed using 2-way ANOVA followed by either Tukey’s multiple comparison tests (Fig A), 1-way ANOVA followed by either Tukey’s multiple comparison tests (Fig. C, D, E) and unpaired t-tests (Fig. F). Data are means ± SEM from three independent experiments. n=3 per group. *p<0.05, **p<0.01, ***p<0.001, ****p<0.0001, ns not significant.

Next, to examine if the improved barrier function was related to increased expression of genes for TJ proteins, we treated Caco-2 cells grown to a differentiated epithelium with 8 μM zinc and 100 nM FICZ either separately or in combination and analysed abundance of mRNA for zona occludens-1 (ZO-1), occludin (OCLN) and claudin-3 (CLD-3). Both 8 μM zinc or 100 nM FICZ independently increased expression of *ZO-1, OCLN* and *CLD-3* in Caco-2 cells, with a stronger effect when given as combination treatment (Fig. 4C). Chelation of the 8 μM zinc added to the medium by an equimolar concentration of the cell permeant zinc chelator, tris(2-pyridylmethyl)amine (TPA)^30^, completely abolished the effects of zinc and FICZ on expression of *ZO-1* and *CLD-3* while diminishing those on expression of *OCLN* (Fig. 4C). Overwhelming the chelating capacity of TPA by addition of another 8 μM zinc (16 μM in total) restored the effect of zinc and FICZ on *ZO-1, OCLN*, and *CLD-3* gene expression showing that the effects of FICZ and changing the zinc concentration of the medium on genes for TJ proteins were dependent on intracellular zinc. The effects of zinc and FICZ on TJ gene expression were confirmed in human ileum organoids treated with 0, 25 or 50 μM zinc added to the medium in presence of 10 μM Ca-EGTA (to reduce background zinc in the medium) with or without FICZ (Fig. 4D). In addition, we found that expression of mucin-2 (*MUC2*) mRNA was increased in the human ileum organoids treated with zinc and FICZ (Fig. 4D), supporting the data from the mouse experiment (Figs. 1C, 2D, Extended data Figs. 1C, 2D, 2F) and previous findings in an *in vitro* goblet cell model^27^. Western blot analysis of ZO-1 and OCLN abundance in Caco-2 cells indicated that expression of both TJ proteins increased after treatment with either zinc or FICZ, but treatment of cells with the combination of zinc and FICZ had a stronger effect (Fig. 4E). Again, the effect of zinc addition to the medium on TJ protein expression appeared to be caused primarily by an increase in intracellular zinc concentration as shown by intracellular zinc chelation using TPA (Fig 4E). The effects of zinc and FICZ on OCLN and MUC2 were identified and quantified by immunocytochemistry in human ileum organoids (Fig. 4F). These results demonstrated that treatment of human intestinal epithelial cells with zinc and FICZ induced expression of mRNA for TJ proteins and *MUC2*, and this translated into increased abundance of the respective proteins, providing a plausible explanation for the enhanced barrier function following the same treatment.

We subsequently investigated the mechanisms by which zinc and AHR ligands might enhance abundance of TJ proteins. The NF-κB pathway has been shown to transcriptionally suppress genes for TJ proteins through NF-κB P65 binding to their promoters^31^. Furthermore, AHR activation has proven effective in attenuating NF-κB activation, mainly through interfering with P65 recruitment to DNA^32^. Zinc can inhibit NF-κB signalling by preventing phosphorylation of P65 and IκBα^33^. We hypothesised that zinc and AHR signalling may converge on and effectively inhibit activation of the NF-κB pathway. We therefore investigated the effects of 8 μM zinc and 100 nM FICZ individually or in combination on the NF-κβ pathway in Caco-2 cells treated with tumour necrosis factor alpha (TNF-α) to stimulate activating serine-536 phosphorylation of P65 and serine-32 phosphorylation of IκBα. Addition of either zinc or FICZ to the medium strongly attenuated TNF-α induced phosphorylation of P65 and IκBα and the combination treatment was the most potent in both cases (Fig 5A). We therefore repeated the measurement of expression of genes encoding TJ proteins in Caco-2 cells after treatment with zinc and/or FICZ, but this time in the presence of a NF-κB blocker, EVP4593 (Extended data Fig. 4A). In the presence 10 μM EVP4593, zinc and/or FICZ had no stimulatory effect on expression of *ZO-1* and *OCLN*, indicating that zinc and FICZ increase expression of mRNA for TJ proteins by alleviating transcriptional inhibition by NF-κB.

**Figure 5:**
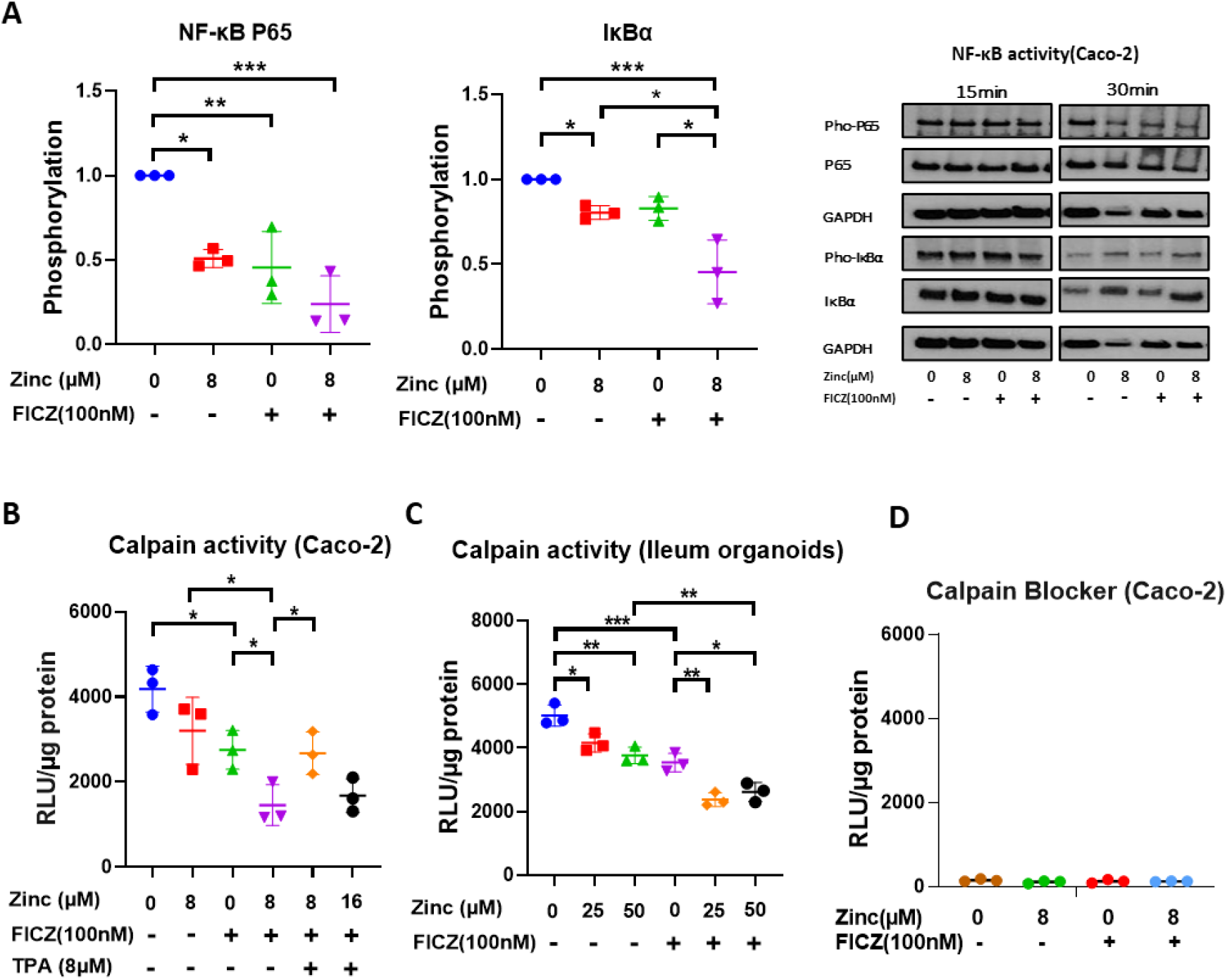
Combined treatment of zinc and AHR agonist FICZ inhibits activities of NF-κß and calpain. (A) Caco-2 cells were treated with TNF-α to stimulate serine-536 phosphorylation of P65 and serine-32 phosphorylation of IκBα in media contain containing 0 or 8 μM zinc with or without 100 nM FICZ after 24 hours. The plots show phosphorylation state of P65 at 30 min and IκBα at 15 min when maximum effect of treatments was observed. (B) Calpain activity in Caco-2 cells treated with zinc and or FICZ for 24 h. Zinc dependence of the effect was shown by addition of a zinc chelator, TPA. (C) Calpain activity in human ileum organoids treated with different concentrations of zinc with or without FICZ for 24 h. (D) Calpain activity in Caco-2 cells after application of 50 μM calpain blocker, calpeptin. Statistical analysis of the data was performed using 1-way ANOVA followed by either Tukey’s multiple comparison tests. Data are means ± SEM from three independent experiments. n=3 per group. *p<0.05, **p<0.01, ***p<0.001, ****p<0.0001, ns not significant.

TJ proteins are degraded by calpain proteases, which are inhibited by zinc^34,35^. We therefore investigated the effects of 8 μM Zinc and 100 nM FICZ individually or in combination on calpain activity in Caco-2 cells grown to an epithelium and in human ileum organoids. Addition of either zinc or FICZ to the medium strongly reduced calpain activity in Caco-2 cells and human ileum organoids, and the combination treatment was the most potent in inhibiting calpain (Fig 5 B, C). We therefore confirmed the expression on TJ proteins after zinc and/or FICZ treatment in Caco-2 cells grown to a differentiated epithelium, but at this time in the presence of a calpain inhibitor, calpeptin (Extended data Fig. 4B). In the presence of 50 μM calpeptin, zinc and/or FICZ had no stimulatory effect on abundance of TJ proteins, suggesting that inhibition of calpain is a second mechanism by which zinc and FICZ improve Caco-2 cell epithelium barrier function.

Because AHR activation by FICZ influenced expression of TJ genes and proteins in a zinc-dependent manner we tested if AHR activation by FICZ induces transcription of genes for zinc regulatory proteins. We previously published, based on chromatin immunoprecipitation microarray (ChIP-chip) mouse data obtained from Dere et al.^36^, that AHR has functional binding sites in several zinc-regulatory genes^37^. Analysis of a more recent ChIP-seq dataset^38^ from human MCF-7 cells exposed to the xenobiotic AHR agonist, 2,3,7,8-tetrachlorodibenzo-p-dioxin (TCDD), confirmed AHR binding to numerous zinc-regulatory genes, including genes for several zinc importers of the ZIP (SLC39) family. To determine if FICZ induces expression of the zinc-regulatory genes identified as potentially being trans-activated by AHR, we measured expression of these genes in differentiated Caco-2 cells as well as in human intestinal organoids following treatment with 10 or 100 nM FICZ. Transcripts for zinc transporters, *ZĨP2, −4, −6, −7* and *-10*, which mediate zinc flux into the cytosol^39^, showed increased mRNA abundance in differentiated Caco-2 cells following treatment with 10 or 100 nM FICZ, but without clear difference between the two concentrations (Fig. 6A). There was also increased expression of metal-responsive transcription factor 1 (*MTF1*), which is an intracellular Zn^2+^ sensor and transcriptional regulator of several zinc homeostatic genes, including metallothionein 1A (*MT-1A*) and *ZIP10*. In human ileum organoids, increased gene expression was observed for *MTF1, ZIP4, −6, −7*, and *-10* (Fig. 6B) following treatment with FICZ; while in wild type mouse ileum organoids there was significantly increased expression of *Zip4, −6, −7*, and *-10* (Fig. 6C). We used organoids derived from ileum of villin^cre^*Ahr*^fl/fl^ mice to investigate if the changes in gene expression were dependent on AHR activation. FICZ was unable to induce expression of any of the genes investigated in the AHR-deficient ileum organoids (Fig. 6D). To establish if FICZ-induced gene expression is associated with AHR recruitment to these genes, we carried out ChIP of AHR in FICZ-treated Caco-2 cells followed by qPCR of the fragments with AHR binding sites in the vicinity of the genes of interest based on previous studies^37,38,40^. We were able to detect statistically significant recruitment of AHR to *ZIP4, ZIP6, ZIP7, ZIP10* and *MTF-1*. Of these genes, *ZIP4* has a binding site for AHR within 1,000 bp of the transcription start site (Extended data Table 1) and is the most likely one to be directly regulated by AHR recruitment to its promoter. ZIP4 is essential for intestinal zinc uptake, the intestinal stem cell niche, and survival. Its transactivation by AHR is therefore of paramount significance. Thus, AHR regulates expression of several zinc regulatory genes in the intestinal epithelium, notably including the gene for ZIP4.

**Figure 6:**
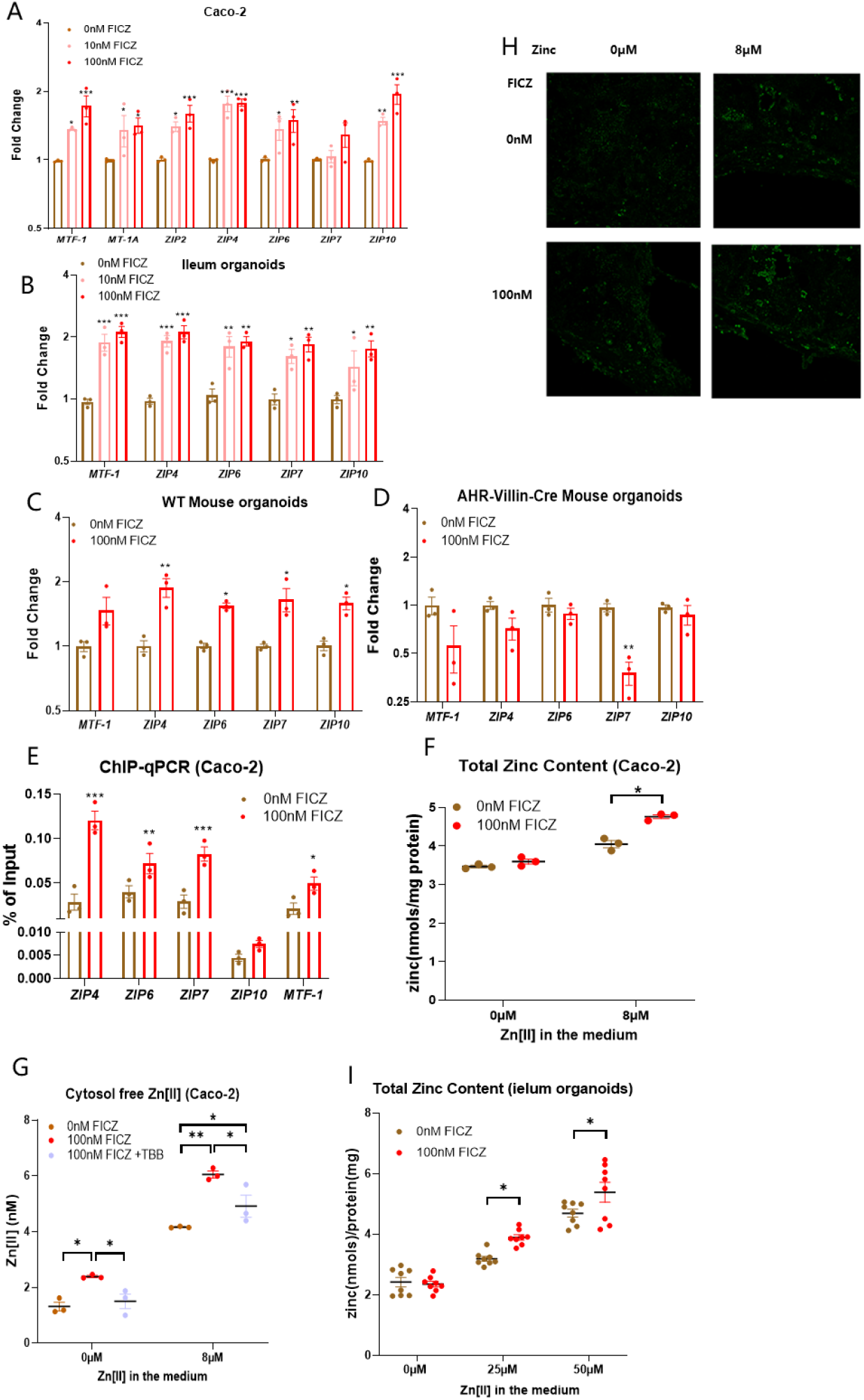
AHR regulates expression of zinc importers and other zinc homeostatic genes and stimulates cellular zinc accumulation. Expression of zinc homeostatic genes in (A) Caco-2 cells, (B) human ileum organoids, (C) ileum organoids from WT mice, and (D) ileum organoids from villin^cre^ *Ahr*^fl/fl^ mice after treatment with FICZ for 24 h. (E) ChIP-qPCR data showing occupancy of AHR on DNA fragments of zinc homeostatic genes from Caco-2 cells treated with FICZ for 24 h. (F) Total zinc content or (G) free cytosolic Zn^2+^ concentrations (measured with FluoZin-3AM probe) in Caco-2 cells following incubation with media containing 0 or 8 μM zinc with or without 100 nM FICZ. A CK2 inhibitor, TBB, was used to prevent activating phosphorylation of ZIP7. (H) Imaging of the fluorescent Zn^2+^ probe, FluoZin-3AM, in Caco-2 cells after 24 h incubation in media containing 0 or 8 μM zinc with or without 100 nM FICZ. (I) Total zinc content in human ileum organoids treated with different concentrations of zinc with or without FICZ for 24 h. Statistical analysis of the data was performed using unpaired t-tests (E) and 2-way ANOVA (A-D, F-I) followed by either Tukey’s or Sidak’s multiple comparison tests. Data are means ± SEM from three independent experiments. n=3 per group. *p<0.05, **p<0.01, ***p<0.001, ****p<0.0001, ns not significant.

To test if AHR-dependent expression of zinc importers translates to increased zinc content and cytosolic free Zn^2+^ concentration we treated differentiated Caco-2 cells with 100 nM FICZ for 24 h in absence or presence of 8 μM extracellular zinc and measured total cellular zinc content by ICP-MS and cytosolic free Zn^2+^ concentrations by the fluorescent Zn^2+^ probe, FluoZin-3AM. In the presence of 8 μM extracellular zinc, treatment with 100 nM FICZ increased the cellular zinc content, but no change was observed when there was only background (<0.0010 μM) zinc in the medium (Fig. 6F, G, H). Similarly, human ileum organoids treated with FICZ in presence of different additions of extracellular zinc (0, 25, 50 μM) showed FICZ-dependent increase in total zinc content when extracellular zinc was available (Fig. 6I). This indicates that FICZ stimulates zinc uptake in differentiated Caco-2 cells and human ileum organoids consistent with the observed upregulation of zinc importers. Whilst the concentration of free cytosolic Zn^2+^ was much lower in cells incubated for 24 h with <0.001 μM zinc in the medium compared with those kept in 8 μM zinc, treatment with FICZ resulted in an increased concentration of cytosolic free Zn^2+^ whether or not zinc was present in the medium (Fig. 6G, H). These results indicate that FICZ treatment can also cause release of Zn^2+^ into the cytosol from intracellular stores. Zinc importers ZIP2, −4, −6, and −10 can mediate cellular zinc uptake^41^, but ZIP7 releases zinc from the endoplasmic reticulum (ER) into the cytosol^41^. Increased expression of *ZIP7* following FICZ treatment as reported above would explain the observation that FICZ treatment results in increased cytosolic free Zn^2+^ in the absence of extracellular zinc. ZIP7 zinc transporting activity is dependent on serine phosphorylation by casein kinase-2 (CK2)^42^. To test whether FICZ increases ZIP7-mediated Zn^2+^ release into the cytosol, tetrabromobenzotriazole (TBB), a selective CK2 blocker, was used to reduce ZIP7 zinc transport activity, and intracellular zinc was monitored by FluoZin-3AM (Fig. 6G). TBB treatment abolished the FICZ-stimulated increase in cytosolic free Zn^2+^ in the absence of extracellular zinc demonstrating that AHR activation does increase ZIP7-dependent flux of zinc from the ER into the cytosol. These experiments allow us to conclude that AHR-dependent expression of ZIP zinc importers results in an increased accumulation of total zinc and cytosolic free Zn^2+^.

In this investigation, we demonstrate that AHR activation promotes barrier function by increasing tight-junction formation and mucus production by mechanisms that involve zinc uptake and increased free cytosolic Zn^2+^. Specifically, we show that activation of AHR by physiologically relevant ligands induces expression of zinc importers of the ZIP family, which facilitate uptake of zinc into the epithelial cells and increases the cytosolic Zn^2+^ concentration. Zn^2+^ is an intracellular signalling ion which participates in a multitude of pathways, including those that regulate the development, function, and integrity of the intestinal epithelium^43–45^. Our data suggest that Zn^2+^ enhances expression of genes for TJ proteins by inhibiting the NF-κB pathway, which otherwise supresses expression of genes for the TJ proteins^46^. We also show that zinc inhibition of calpain activity helps to maintain the levels of TJ proteins.

It has been shown previously that OCLN expression can be enhanced following stimulation of the Zinc Receptor, G-coupled Protein Receptor-39 (GPR39), with extracellular zinc^47^. However, AHR activation did not affect *GPR39* expression in mouse ileum and colon, or human ileum organoids (Extended data Fig. 4C, D) and it is therefore unlikely that the AHR-stimulated zinc-dependent beneficial effects on epithelial integrity involved activation of GPR39 by extracellular zinc.

In this study, AHR-stimulated zinc uptake, reduced epithelial leakiness of differentiated Caco-2 cells and human ileum organoids, and alleviated DSS-induced IBD in ileum and colon of mice. The therapeutic efficacy of I3C was completely lost in mice that had intestinal epithelial deficiency of AHR showing that the effect was mediated by AHR specifically in these cells. Moreover, whilst in wild-type mice zinc had beneficial effects on IBD model symptoms, in mice lacking the AHR in the intestinal epithelium the DSS-induced lesions were equally severe in the animals receiving a zinc-supplemented diet (100 mg/kg feed) as in those given zinc-depleted feed (5 mg/kg). It is also worth noting that the efficacy of combined zinc and AHR agonist treatment was more pronounced in the animal model than in our *in vitro* experiments. Whereas in wild-type mice combined treatment with I3C and zinc almost completely prevented DSS-induced injury, the effect in the Caco-2 cell injury models was partial. Both AHR and zinc have a multitude of effects on the gastrointestinal tract notably including effects on the gastroenteric immune system, which can explain the higher efficacy *in vivo* than *in vitro*^26,48,49^.

Our findings conclusively show that physiological dietary AHR agonists are efficacious in preventing and perhaps treating IBD, but only in combination with sufficient zinc intake because much of the therapeutic effects of AHR are zinc dependent. This could be critically important because approximately 1/3 of the World’s population is zinc-deficient with the prevalence being highest in low-income countries where diets are dominated by plant-based foods^50^. Whilst plant-based foods are the main source of AHR pre-ligands they also contain phytate, which makes zinc unavailable for uptake^51^. Meat and seafood are generally considered the best dietary sources of zinc, but sufficient zinc intake in absence of these food groups could be solved with dietary supplementation or by fortifying other foods with zinc^52^. Such strategies may also be required in industrialised countries to combat a drift toward lower dietary zinc availability driven by global sustainability issues which are prompting a move away from animal-based diets, toward plant-based foods.

A major fundamental discovery in the present study is that AHR is an important regulator of zinc fluxes and zinc-dependent processes in the intestinal epithelium. Not only does AHR regulate expression of zinc transporters, sensors and binding proteins, but in the absence of AHR the therapeutic effect of zinc on DSS-induced IBD is completely lost (Figure 2; Extended data Figure 2). This novel finding is likely to have wider implications on other physiological and pathophysiological processes throughout the body as zinc transporters and Zn^2+^ ions participate in regulation of virtually every aspect of biology including fertilisation, cell division, differentiation, growth, immunity, and cognition^53,54^.

## Authors’ Contributions

Xiuchuan Hu (XH) performed and analysed most of the experiments in cell line and organoids with input from Xinyi Li, Gavin A. Bewick provided human ileum organoids with organoids isolation and culture by Yuxian Lei and Muralidhara Rao Maradana provided mice ileum organoids. Christer Hogstrand (CH), Peter Kille (PK) and Wolfgang Maret (WM) conceived and designed the study with input from GB, XH and Yufeng Zhou (YZ). Wenfeng Xiao (WX) performed and analysed most mice experiments with assistance by Yaojing Gao and Xueru Xie., M. Albert Basson provided expertise and support for ChIP and mouse experiments. CH and YZ drafted the manuscript together with XH and WX. All authors contributed to editing of the manuscript.

## Acknowledgments

The authors wish to thank Dr Brigitta Stockinger at the The Francis Crick Institute, London, UK, for valuable advice on the experiments and the manuscript. This work was supported by grants from Guts.UK (DGN2019_02 to CH), ZinPro Performance Minerals (1117612 to CH), from the National Key R&D Program of China (2021YFC2701800, 2021YFC2701802 to YZ), National Natural Science Foundation of China (81974248 to YZ), Program for Outstanding Medical Academic Leader (2019LJ19 to YZ), and Shanghai Committee of Science and Technology (21140902400 to YZ). XH was supported by a studentship from Chinese Scholarship Council and King’s College London. Metal analysis was performed by the London Metallomics Facility, which was funded by the Wellcome Trust (202902/Z/16/Z to WM). Analysis of gene expression by qPCR was carried out in King’s College London Genomics Centre.

## Methods

### Mice

All WT and genetically modified mice used were on the C57BL/6J background. *Ahr* floxed (B6.129(FVB)-*Ahr^tm3.1Bra/J^*) mice were purchased from the Jackson Laboratory (Bar Harbor, ME, USA). Vil1-cre mice [B6.Cg-Tg 1000 Gum] were obtained from Shanghai Model Organisms (Shanghai, China). *Ahr* floxed mice and Vil1-Cre mice were crossed for obtaining specific deletion of *Ahr* in villus epithelial cells of the small and large intestines (villin^cre^*Ahr*^fl/fl^). C57BL/6J mice were purchased from SLAC laboratory animal (Shanghai, China). All mice were housed, bred, and maintained under specific pathogen-free conditions. All experiments complied with the relevant laws and institutional guidelines, as overseen by the Animal Studies Committee of the Children’s Hospital of Fudan University.

### DSS-induced IBD model

For WT mouse experiments, three-week-old female C57BL/6J mice were randomly divided into 12 groups based on following treatments alone and in combinations: zinc content (5, 35, 100 mg/kg) diet, I3C or ddH2O (by oral gavage) as control, with tap water for drinking with or without 2% DSS. Zinc diets (SYSE BIO, Changzhou, China) and water with or without DSS (MP Biomedicals, OH, USA) were given *ad libitum*. I3C (Sigma, Milwaukee, WI, USA) was given at a dose of 1 mg/mouse dissolved in 100 μl ddH2O by daily gavage. Zinc diets and I3C were given from the beginning to the end of the experiment and DSS was given from Day 4 to Day 11 when mice were sacrificed by terminal anaesthesia. For the villin^cre^*Ahr*^fl/fl^ mouse experiment, 3-4 weeks old female mice were randomly divided into the same 12 groups as in the experiment with WT mice. The different zinc diets and I3C were given from beginning to end of the experiment and DSS was given from Day 4 to Day 10. Mice were sacrificed on Day 10 (not on Day 11) because of animal welfare due to serious colitis symptoms.

### Determination of disease activity index

Body weight, gross blood, and stool consistency were analysed daily. Disease activity scores were analysed according to previously published methods^56^. Briefly, for stool consistency, 0 points were assigned for well-formed pellets, 2 points for pasty and semi-formed stools that did not adhere to the anus, and 4 points for liquid stools that did adhere to the anus. For bleeding, 0 was assigned for no blood, 1 point for visible slight bleeding, and 4 points for gross bleeding.

### Histology

Colons and ileum were fixed in 10% neutral-buffered formalin. Paraffin sections were stained with hematoxylin and eosin (H&E) and Periodic Acid-Schiff ^55^. Colon histological scoring was determined by the combination of scores for inflammatory cell infiltration (score, 0 to 3) and tissue damage (score, 0 to 3) as published previously^30^. Regarding scores for inflammatory cell infiltration, 0 points were assigned for the presence of occasional inflammatory cells in the lamina propria, 1 point for increased numbers of inflammatory cells in the lamina propria, 2 points for confluence of inflammatory cells extending into the submucosa, and 3 points for transmural extension of the infiltrate. For scoring of tissue damage, 0 points were given for no mucosal damage, 1 point for lymphoepithelial lesions, 2 points for surface mucosal erosion or focal ulceration, and 3 points for extensive mucosal damage and extension into deeper structures of the bowel wall^56^.

Ileum histological scoring was determined by inflammatory cell infiltrate, epithelial changes and mucosal architecture together as follows: 0 points were assigned for normal ileum with intact epithelium and short, finger like villi, 1 point for mild mucosal inflammatory cell infiltrate, 2 points for mild diffuse inflammatory cell infiltrate in mucosa and submucosa, 3 points for moderate inflammatory cell infiltrates in mucosa and submucosa with villous blunting, 4 points for marked mucosal, submucosal and transmural inflammatory cell infiltration accompanied by villous broadening and 5 points for marked transmural inflammatory cell infiltration and villous atrophy^57^.

### Immunohistochemical staining and scoring

For immunohistochemistry (IHC), 6 μm frozen colon sections were incubated with primary antibodies against MUC2 or Occludin, and then incubated with secondary antibodies, followed by staining and imaging. Details of the antibodies used are provided in the Extended data Table 3. Six fields under 400x magnification were selected from each group in a blinded way. The expression level was scored using the IHC profiler^58^ plugin in ImageJ software, which allows for non-biased, automated scoring of histological slides. In this method, each DAB-stained pixel is categorized into one of four pre-set pixel intensity bins (High Positive, Positive, Low Positive, Negative). And 0 points were assigned for negative, 1 for low positive, 2 for positive and 3 for high positive.

### Microbiome profiling

The colon contents of mice were used for Microbiome profiling. Total bacterial DNA was extracted by the manufacturer’s protocol (the Power Soil DNA Isolation Kit, MO BIO Laboratories). OD260/280 and OD230/260 were measured to assess the quality and concentration of extracted total DNA. The 16S rRNA genes were subjected to polymerase chain reaction amplification using the universal primer (Forward primer, GTGAATCATCGARTC; reverse primer, TCCTCCGCTTATTGAT) targeting V3 + V4 regions. High-throughput sequencing analysis of bacterial 16S rRNA genes was performed based on the Illumina Novaseq at Biomarker Technologies Corporation (Beijing, China). According to the relationship between the overlap and paired-end reads, raw tags were obtained from the splicing sequence using Trimmomatic (version 0.33). Then, Cutadapt (version 1.9.1) was used to obtain clean tags and effective tags, respectively. Clustering of the effective tags into operational taxonomic units (OTUs) at 97% similarity was performed using USEARCH (version 10.0). Then, OTUs were annotated using RDP Classifier (version 2.2, at 0.8 confidence threshold) based on the Silva taxonomy database. Finally, alpha diversity, beta diversity and phenotype prediction were performed using QIIME2, QIIME and Bugbase, respectively.

### Generation and culture of mouse ileum organoids

Mouse ileum crypts were isolated from the ileum segment of WT and villin^cre^*Ahr*^fl/fl^ mice in a PBS solution containing 2 mM EDTA. The tissue was incubated for 30-45 mins in a 4 °C shaking incubator (200RPM), followed by washing and manual shaking in cold D-PBS to isolate crypts. To generate organoids, isolated crypts were embedded in Matrigel and resulting organoids were maintained in cell growth medium (WENR) supplemented with 10 μM Y-27632 (Sigma–Aldrich) was overlaid. WENR medium consists of advanced Dulbecco modified Eagle medium (DMEM)/F12, 1X GlutaMAX, 10 mM HEPES, 100 units/mL penicillin/streptomycin, 50 ng/mL EGF, 1× B27, 1× N2 supplements (all from Gibco), 1 mM N-acetylcysteine (Sigma–Aldrich), 100 ng/mL Mouse Noggin (Peprotech) and 10% R-spondin-1 conditioned medium (lab production). Three days later, the medium was changed into differentiation media ^59^ with no Y-27632. Organoids were passaged once a week by mechanical dissociation, at a 1:3 split ratio. Plated organoids were maintained at 37°C in an incubator with 5% CO_2_, and the media were changed every other day.

### Generation and culture of human terminal ileal organoids

Preparation of human terminal ileal organoids has been described before^60^. Briefly, human terminal ileum crypts were isolated from biopsies acquired from patients undergoing colonoscopy at Guy’s and St. Thomas’s NHS Foundation Trust with their informed consent. Biopsies were washed in cold PBS until the supernatant was clear. Following 10 min of incubation at room temperature with 10 mM 1,4-dithiothreitol (DTT), the biopsies were incubated with 8 mM EDTA in PBS and placed in a rotator for 1 h at 4 °C. At the end of the incubation, the EDTA was removed, and crypts were released with vigorous shaking in cold PBS. The crypts were further washed in PBS, pelleted and resuspended in Matrigel (Corning), in the same density as mouse crypts. Human intestinal crypts embedded in Matrigel were overlaid with stem cell growth medium (WENRAS) supplemented with 10 μM Y-27632 and 5 μM CHIR99021 (Sigma–Aldrich). The human stem cell growth medium, in addition to the components of the previously described mouse medium, also contained 10 nM gastrin (Sigma–Aldrich), 500 nM A83-01 (Bio-techne), 10 μM SB202190 (Sigma–Aldrich) and 10 mM nicotinamide (Sigma–Aldrich). Three days after isolation or splitting, Y-27632 and CHIR99221 were removed from the medium, and organoids were either maintained in WENRAS or transferred into differentiation medium for setting up experiments. For the differentiation of human ileal organoids, Wnt3A surrogate protein in the medium was reduced from 0.15 nM to 0.045 nM and SB202190 and nicotinamide were withdrawn from the medium. Differentiation medium was used for 4 days, and human terminal ileal organoids were treated with different concentration of zinc with or without FICZ for 24 h. Zinc content in differentiated medium and WENRAS was measured by inductively coupled plasma mass spectroscopy (ICP-MS) and determined as 25 μM. A zinc depleted medium was created by addition of 10 μM CaEGTA to WENRAS. Addition of 10 μM CaEGTA together with 25 μM or 50 μM were considered to represent replete zinc supply and zinc supplementation, respectively.

### Cell Culture

Human epithelial colorectal adherent Caco-2 cells, a gift from Prof. Paul Sharp, King’s College London, were maintained in Minimum Essential Medium Eagle (MEM, Merck), supplemented with 10% fetal bovine serum (FBS, Merck), 1% 100× MEM Non-essential Amino Acid Solution (Merck), 1% 250 μg/ml Amphotericin B (Merck), and 1% 100 units/ml penicillin (Merck), at 37°C in a humidified atmosphere of 95% air and 5% CO_2_. The cells were grown in the inserts of 24-well plate for more than 2 weeks at which point they differentiated into an epithelium with properties similar to enterocytes in small intestine as described previously^29^.

### Whole mount staining of organoids

Organoids were isolated from the Matrigel using wash medium (advanced DMEM/F12, 10 mM HEPES and 1X GlutaMAX), and then fixed for 45 mins in 4% paraformaldehyde at room temperature. After washing using PBS-B (PBS, 0.1% BSA), organoids were permeabilized and blocked by incubation in PBS containing 3% BSA, 0.3% Triton-X-100, 1% DMSO and 5% normal goat serum (Merck) for 1 h at RT and incubated overnight with primary antibodies. The next day, organoids were washed with PBS-B and incubated 2 h at room temperature with secondary antibodies. After washing, organoids were mounted with VECTASHIELD Vibrance Antifade Mounting Medium (Vector Laboratories). Images were captured on the Nikon A1 inverted confocal microscope using Z-stacks and processed with ImageJ.

### RNA extraction and quantitative real-time PCR

Total RNA of Caco-2 cells and human ileum organoids was extracted using RNAdvance Tissue kit (Beckman Coulter) and total RNA of mouse organoids was isolated using Quick-RNA Miniprep Plus Kit (Zymo Research), and all of them were assessed for purity and quantity using a Nanodrop 1000 spectrophotometer. High-Capacity RNA-to-cDNA Kit (Fisher Scientific) was used for cDNA synthesis. qPCR assays were designed using the online Universal Probe Library (UPL) assay design tool (https://lifescience.roche.com/en_gb/brands/universal-probe-library.html). Assay designs are provided in Extended data Table 2. PCR plates were loaded using the Biomek FX liquid handling robot (Beckman Coulter) and reactions [10 ng cDNA, 0.1 μM UPL probe, 0.2 μM forward primer, 0.2 μM reverse primer and 5 μL Luna^®^ Universal Probe qPCR Master Mix (New England Biolabs)] amplified using the Prism7900HT sequence detection system (Applied Biosystems) and analysed using sequence detection systems v2.4 software. Human or mouse Glyceraldehyde-3-Phosphate Dehydrogenase (GAPDH), beta-actin (β-actin) or ubiquitin C (UBC) (TableS3) were used as housekeeping genes.

### The transepithelial electricity resistance (TEER) monitor and permeability assay

Caco-2 cells (1 ×10^5^) were grown on 0.336 cm^2^ Transwell inserts (Greiner Bio-One). TEER was measured with was monitored daily using an epithelial tissue volt ohmmeter (EVOMX) with STX-2 chopsticks (World Precision Instruments). TEER measurements were calculated in ohms cm^2^ after subtracting the blank value for the membrane insert. Same inserts were applied for intestinal permeability assay. FITC-dextran 4000 (Merck), 1 mg/ml solution was put in the apical and 100 μL basement medium at 4, 8, 12, and 24 h was measured by fluorometry (excitation, 475 nm: emission, 530 nm). Serial dilutions of FITC-dextran in medium were used to calculate a standard curve.

DSS was used to generate an inflammation model as described before^61^. Briefly, 2% DSS was added in the apical and basement. Zinc and FICZ treatment, and TEER and permeability measurements were the same as normal condition.

Caco-2 cell epithelia were subjected to a hypoxia challenge as described before^62^. Briefly, cells grown in 0.336 cm^2^ Transwell inserts were transferred from a standard CO_2_ incubator with approximately 18% O_2_ to an atmosphere-regulated workstation set to 0% O_2_ and 5% CO_2_ (Sci-tive;Baker-Ruskinn, Sanford, ME, USA) for 6 hours. Media were changed in the workstation to media pre-equilibrated to 0% O_2_. After the 6-h hypoxia challenge, cells were returned to the standard CO_2_ incubator and treated with zinc and/or FICZ. TEER and permeability measurements were the same as normal condition.

Treated human ileum organoids were washed twice with warmed PBS and incubated with 4 kDa FITC-dextran at a final concentration of 1.25 μM at room temperature for 1 hour to impose a chemical serosa-to-lumen gradient. Afterwards, the FITC-dextran was removed, and the organoids were gently washed three times with PBS to remove FITC-dextran from the medium. Domes were resuspended in PBS in Eppendorf tube and washed twice, before using P1000 to seed in the Petri dish. Images were captured on a BioStation IM-Q (Nikon) and fluorescence within the organoids determined (ImageJ) by focusing on the entire luminal area of the organoid.

### Chromatin immunoprecipitation (ChIP)-qPCR

Caco-cells (8×10^6^) grown on 10 cm^2^ dishes were fixed with 1% formaldehyde, and the cross-linking reaction was stopped by addition of 1M glycine. After cell lysis and isolation of nuclei, samples were sonicated in a Bioruptor UCD-300 in 10 mM TRIS pH 8, 1 mM EDTA, 0.5 mM EGTA, 0.5% N-lauroylsarcosine to 200–500 bp fragment size. For each ChIP, 100 μg of chromatin was used, and 1/50 of chromatin as INPUT. Chromatin was immunoprecipitated with 1.1 μg of a rabbit IgG against AHR (83200S, Cell Signaling Technology) and 1.1 μg Normal Rabbit IgG (2729S, Cell Signaling Technology). Complexes were captured with Protein G Dynabeads, washed with modified RIPA buffer (50 mM HEPES pH7.5, 1 mM EDTA, 0.3% Sodium deoxycholate, 1% NP40, 250 mM LiCl), eluted in 50 mM TRIS pH8, 10 mM EDTA, 1% SDS, cross-links reversed by overnight incubation at 65°C and DNA precipitated after phenol-chloroform extraction. AHR-target zinc regulatory genes were identified in ChIP-Atlas^63^ and peaks in these genes visualised using the Integrative Genomics Viewer^64^. DNA sequences corresponding to peaks were downloaded from Ensembl (GRCh37/hg19) and interrogated for AHR elements (AHRE) by the PROMO online tool^65^, which used version 8.3 of the TRANSFAC database. Unique DNA fragments were amplified and quantified by qPCR with primers directed towards AHR-binding loci containing identified AHRE in *MTF1, ZIP4, ZIP6, ZIP7* and *ZIP10* (Extended data Table 1). Relative ChIP enrichment was calculated by 2% input.

### Total zinc analysis

Caco-2 cells or human ileum organoids were lysed in 1 mL 20 mM Suprapur NaOH (VWR International) for 2 hours. The liquid was evaporated to dryness using Genevac™ miVac Quattro Concentrator, then dissolved with 400 μL trace element grade 65-69% HNO_3_ (Merck) and 100 μL H_2_O_2_ (Merck) and digested with lid overnight at 60°C. The samples were then diluted by adding 6 mL Ultrapure 18.2 MOhms water. The zinc concentrations were determined through ICP-MS carried out using a NexIon 350 D (Perkin Elmer) at the London Metallomics Facility. The total cellular zinc content was normalised to protein content.

### Imaging of intracellular Zn^2+^

After incubation with 1 μM FluoZin-3AM for 30 minutes at room temperature, cells were then washed and incubated in HHBSS for another 15 minutes at room temperature before any imaging and readings were taken. Fluorescence was measured with 492 nm excitation and 517 nm emission in a fluorescence plate reader. Pyrithione (5 μM) together with 20 μM ZnCl2 were used to treat cells for 15 minutes for maximum fluorescent reading, and 100 μM TPEN (N, N, N’, N’-tetrakis (2-pyridylmethyl) ethylenediamine) for 15 minutes as minimum fluorescence reading.

### BCA assay

Protein concentrations were measured by the BCA assay. Samples were diluted 1:5 in RIPA buffer, and then 1:50 with BCA assay reagent (Pierce™ BCA Protein Assay Kit, Thermo Fisher Scientific) in a 96-microtiter plate in triplicate. Absorbance was measured at wavelength 595 nm (reference: 450 nm) using a microplate reader and compared to a standard curve generated from dilutions of a known BSA standard (0.125, 0.25, 0.5, 0.75, 1.0, 1.5 and 2.0 mg/ml) (Thermo Fisher Scientific).

### Protein extraction and Immunoblotting

Total protein was extracted using RIPA Lysis supplement with Halt Protease Inhibitor Cocktail and Halt Phosphatase Inhibitor Cocktail (all from Thermo Fisher Scientific) following the manufacturer’s protocol. Briefly, after being washed by cold PBS, cells were scraped and centrifuged for 10 minutes at 15,600 *g* at 4°C and the protein lysate aspirated. Protein concentration was measured by BSA assay as described above and 20 μg Lysates were separated SDS-PAGE Gel. Proteins were transferred onto 0.45 μm pore size nitrocellulose membranes (GE Healthcare Life Sciences] using NuPAGE™ Transfer Buffer, and membranes blocked through incubation in 5% (w/v) BSA or skim milk (1 h, RT). Membranes were incubated with primary antibodies overnight, followed by HRP-linked secondary antibodies. Details of the antibodies used are provided in the Extended data Table 3. Membranes were washed three times with TBS-T between each step. Membranes were treated with ECL Western Blotting Detection Reagent (GE Healthcare Life Sciences) and visualized on X-ray film (GE Healthcare Life Sciences) using a film imager. Immunoblots were washed with TBST three times, incubated in stripping, and further washed with TBS-T. Membranes were blocked with 5% (w/v) skim milk (1 h, RT) before additional antibody incubation, as described in the above. Relative abundance of proteins was determined by densiometric analysis of the X-ray films using ImageJ, followed by normalisation to GAPDH expression.

### NF-κB activity

NF-κB activity was determined by Western Blot and densitometric analysis of phosphorylated IκBα^ser32/36^ and P65^ser536^ compared with total IκBα and P65, respectively. The phosphorylation state of the proteins was calculated by dividing the densities of the phosphorylated protein band with those of the total protein and normalising to the phosphorylation ratios of the control group.

EVP4593 (QNZ) was used as NF-κB pathway inhibitor ^66^ and a 4-h pre-incubation with 10 μM EVP4593 was used to block NF-κB activity.

### Calpain Activity

Calpain activity was measured in Caco-2 cell or organoids extracts by Calpain-Glo™ Protease Assay (Promega) following the manufacturer’s instructions. In short, cells were lysed in cell lysis buffer (Cell Signaling Technology) and centrifuged for 10 min at 15,000 x *g* at 4°C. The supernatant was collected for the assay. 50 μL of 50 ng/μL protein and 50 μL of the Calpain-Glo reagents were added to the luminescent calpain substrate Suc-LLVY-aminoluciferin in the presence of 1 mM CaCl2. Luminescence was detected by BMG CLARIOstar. Calpeptin has been shown to be an effective calpain blocker^67^. Pretreatment for 4 h with 50 μM Calpeptin was used to block calpeptin activity.

### Statistical analyses

The n values represent individual mice for in vivo experiments and biological replicates in cell line and organoids. Two independent experiments were repeated in animal experiments and three independent experiments were repeated in in vitro assays. Unless otherwise stated, all statistical analyses were carried out through unpaired t-tests, 1-way ANOVA or 2-way ANOVA followed by either Tukey’s or Sidak’s multiple comparison tests, as appropriate. Data were considered statistically significant when p <0.05 and presented as mean ± SEM.

