## Extended Data for "Aryl hydrocarbon receptor utilises cellular zinc signals to maintain the gut epithelial barrier"

Xiuchuan Hu[1#], Wenfeng Xiao[2,3#], Yuxian Lei[4], Adam Green[1], Xinyi Lee[1], Muralidhara Rao Maradana[5], Yajing Gao[2,3], Xueru Xie[2,3], Rui Wang[1], George Chennell[6], M. Albert Basson[7], Pete Kille[8], Wolfgang Maret[1], Gavin A. Bewick[4], Yufeng Zhou[2,3]*, Christer Hogstrand[1]*

[1] Department of Nutritional Sciences, King's College London, London, UK

[2] Institute of Pediatrics, Children's Hospital of Fudan University, and the Shanghai Key Laboratory of Medical Epigenetics, International Co-laboratory of Medical Epigenetics and Metabolism, Ministry of Science and Technology, Institutes of Biomedical Sciences, Fudan University, Shanghai, China.

[3] National Health Commission (NHC) Key Laboratory of Neonatal Diseases, Fudan University, Shanghai, China.

[4] Department of Diabetes, Cardiovascular and Metabolic Medicine & Sciences, Faculty of Life Science and Medicine, King's College London, London, UK.

[5] The Francis Crick Institute, London, UK.

[6] Clinical Neuroscience Department, King's College London, London, UK.

[7]Centre for Craniofacial and Regenerative Biology and MRC Centre for Neurodevelopmental Disorders, King's College London, London, UK

[8] School of Biosciences, Cardiff University, Cardiff, UK

[#] These authors contributed equally to this work.

### Extended data Figure 1

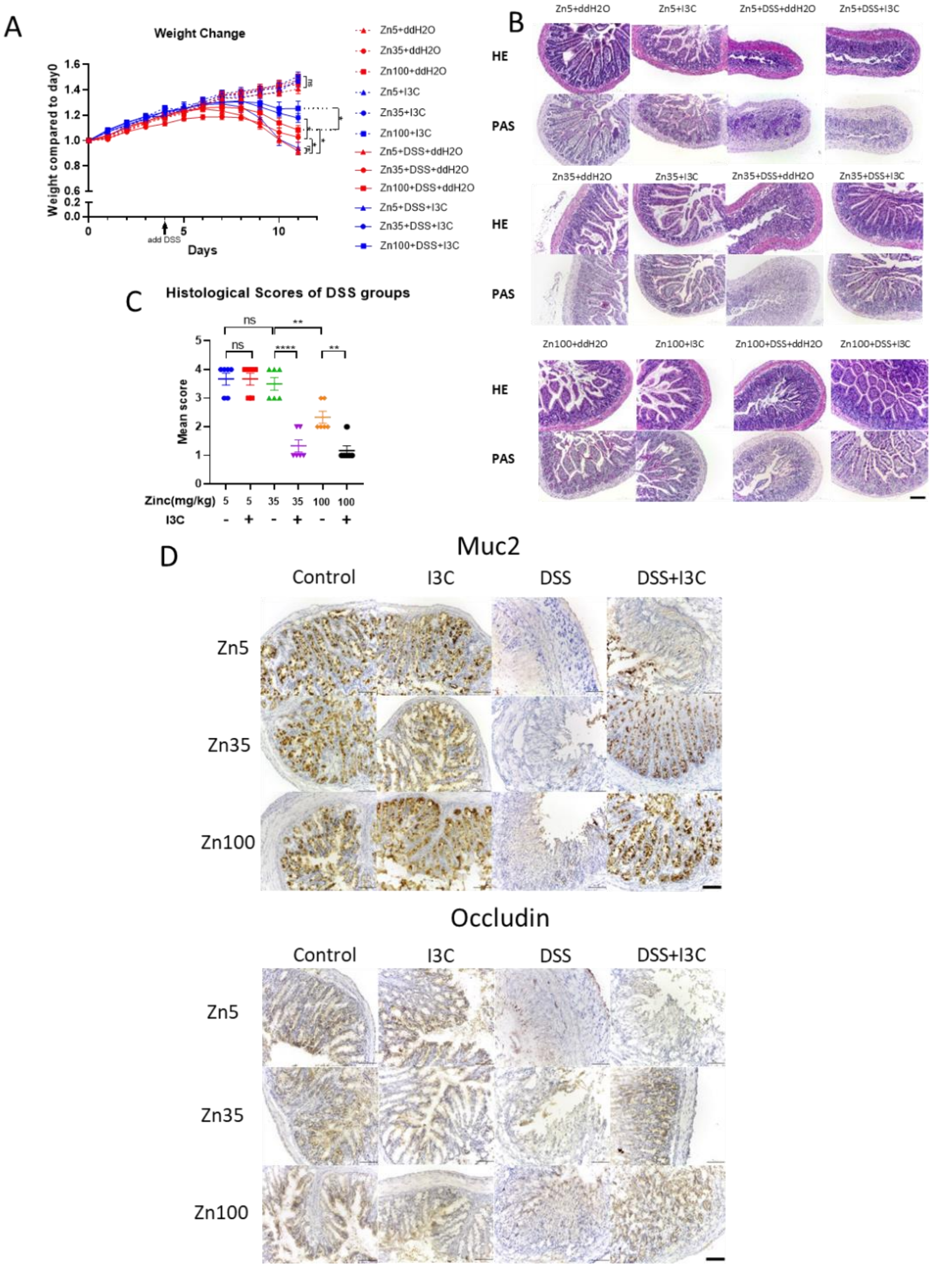


**Extended data Figure 1: Effects of zinc deficiency and I3C treatment in DSS-induced IBD mouse model on body weight, histopathology of ileum, and expression of marker for epithelial barrier function**

Three-week-old C57BL/6J mice were provided with diets with one of three zinc concentrations (5 mg/kg (Zinc5), 35 mg/kg (Zinc35) and 100 mg/kg (Zinc100) from Days 0-10 with (I3C) or without (ddH$_2$O) I3C given by daily gavage. DSS was administered by the drinking water from Day 4 to Day 10. Mice were sacrificed on Day 11. (A) Mean daily weight change compared to Day 0. (B) Histopathological changes in the ileum tissue examined by H&E and Periodic Acid Schiff[55] staining (magnification, ×200). Scale bar, 200 μm. (C) Histopathological scores of the ileum tissue in DSS treated mice. (D) Representative IHC images of MUC2 and Occludin expression in colon tissues (magnification, ×200). Scale bar, 200 μm. Representative data are means ± SEM and n=6 in each group. Animal treatments were repeated twice. Statistical analysis of the data was performed using 1-way ANOVA followed by Tukey's multiple comparison tests. *p<0.05, **p<0.01, ***p<0.001, ****p<0.0001， ns not significant.

### Extended data Figure 2

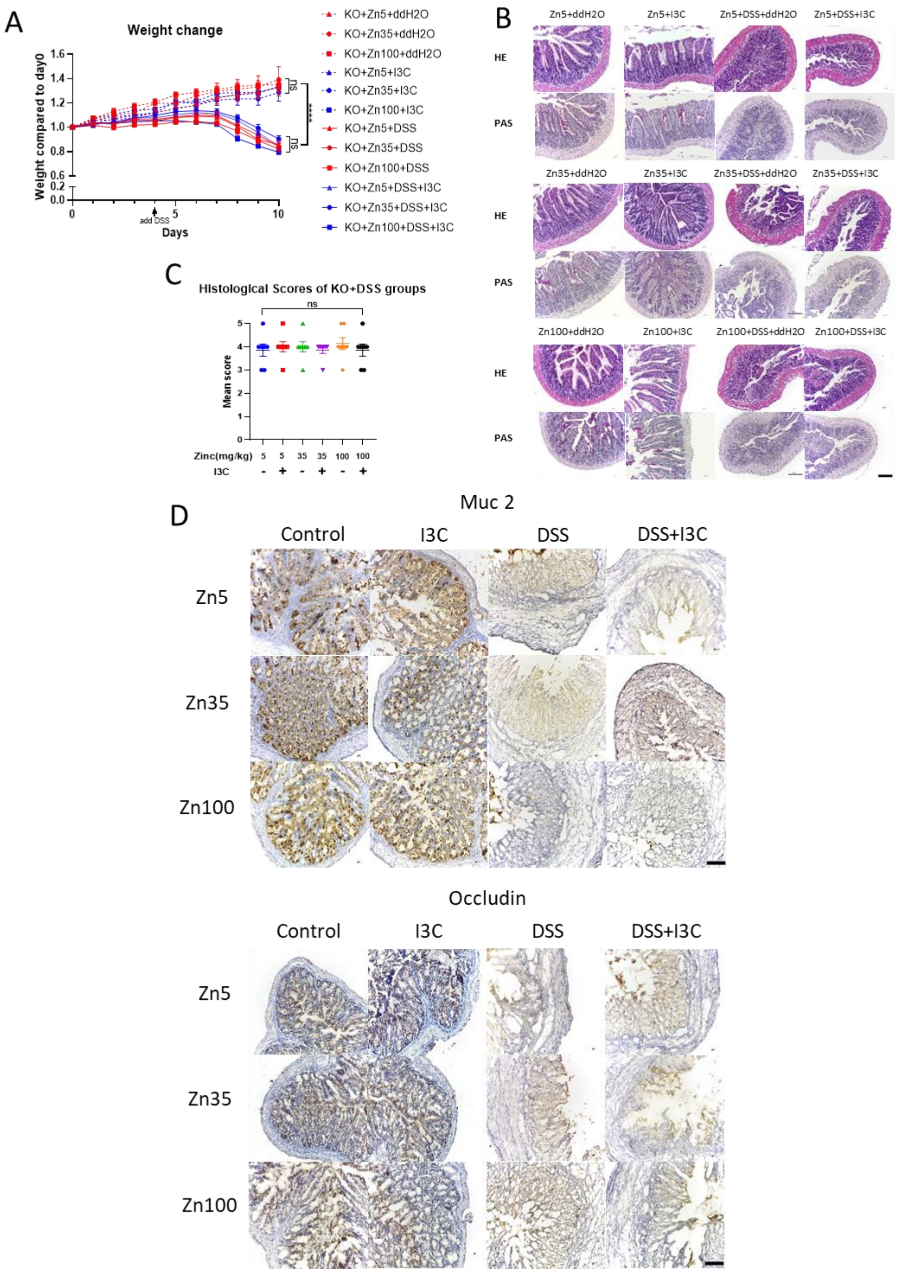


**Extended data Figure 2: Effects of zinc deficiency and I3C treatment in DSS-induced IBD model in Vil1-Ahr KO mice on body weight, histopathology of ileum, and expression of marker for epithelial barrier function**

Three-week-old villin$^{cre}$ *Ahr*$^{fl/fl}$ mice (KO) on C57BL/6J background were provided with diets with one of three zinc concentrations (5 mg/kg (Zinc5), 35 mg/kg (Zinc35) and 100 mg/kg (Zinc100) from Days 0-10 with (I3C) or without (ddH$_2$O) I3C given by daily gavage. DSS was administered by the drinking water from Day 4 to Day10. Mice were sacrificed on Day 11. (A) Mean daily weight change compared to Day 0. (B) Histopathological changes in the ileum tissue examined by H&E and Periodic Acid Schiff[55] staining (magnification, ×200). Scale bar, 200 μm. (C) Histopathological scores of the ileum tissue in DSS treated mice. (D) Representative IHC images of MUC2 and Occludin expression in colon tissues (magnification, ×200). Scale bar, 200 μm. Representative data are means ± SEM and n=6 in each group. Animal treatments were repeated twice. Statistical analysis of the data was performed using 1-way ANOVA followed by Tukey's multiple comparison tests. *p<0.05, **p<0.01, ***p<0.001, ****p<0.0001, ns not significant.

### Extended data Figure 3

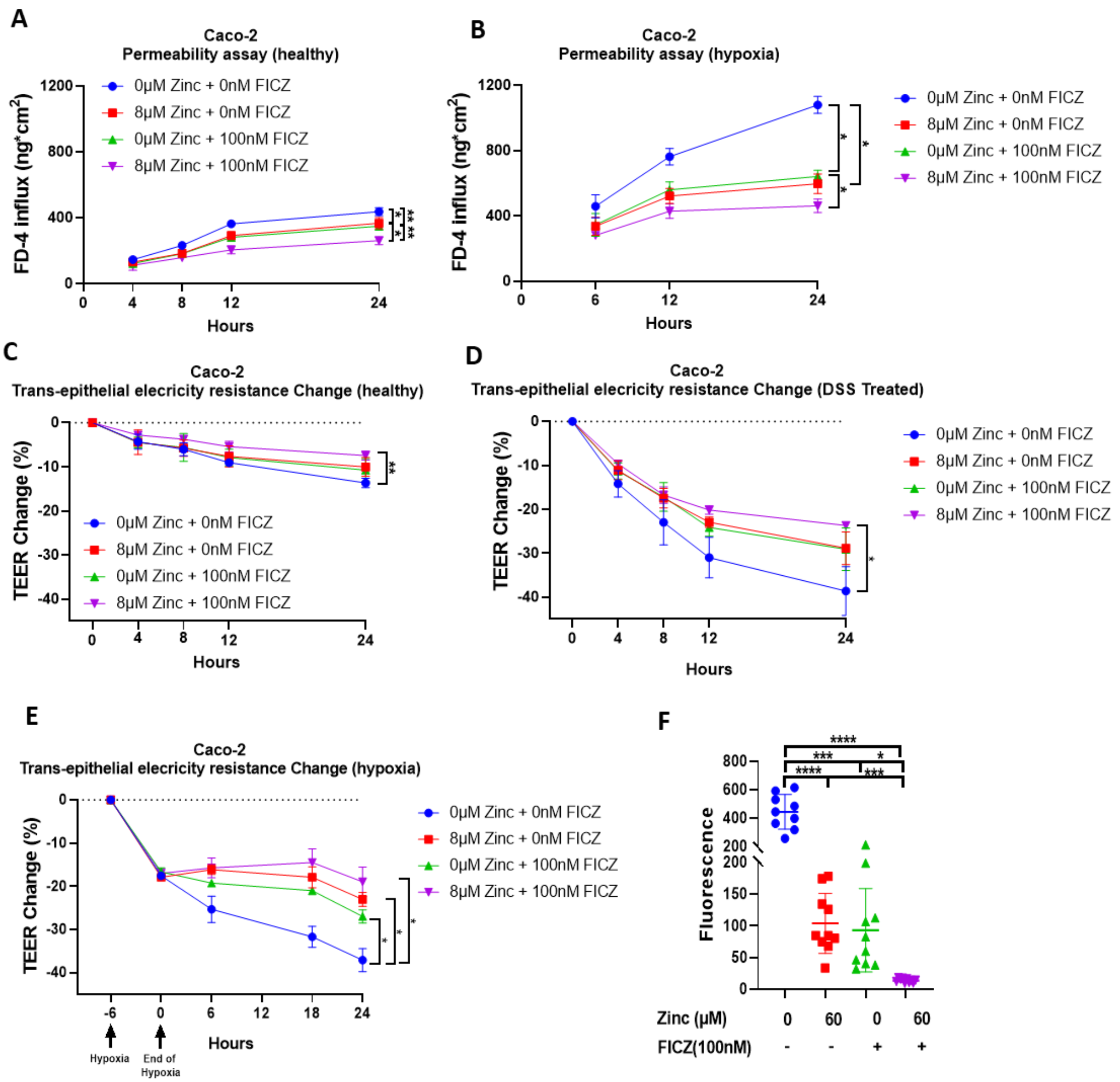


**Extended Data Figure 3: Additional evidence that combined treatment of AHR agonist FICZ and zinc promotes epithelial barrier function in Caco-2 cells and human ileum organoids**

(A-E) Caco-2 cells were grown to an epithelium in a Transwell® system. At the start of the experiment, the medium in the apical compartment was replaced with MEM containing 0 or 8 µM zinc with or without 100 nM FICZ as indicated in the figure. (A,B) FITC-dextran 4000 (Merck) was added to the apical medium at t=0 and permeability was measured by sampling the medium in the basal compartment and measurement of FITC fluorescence over 24 hours. (A) The cells were left unchallenged to represent a healthy gut epithelium or (B) challenged with hypoxia for six hours prior to the start of the experiment and returned to atmospheric $P_{O2}$ at t=0. (C-E) Changes in electrical resistance across Caco-2 cell epithelium kept in media with 0 or 8µM zinc with or without 100 nM FICZ. (A) The Caco-2 cell epithelium was unchallenged during the 24 h period representing a "Healthy" gut epithelium or challenged with (D) 3% DSS or (E) hypoxia for 6 h prior to the addition of zinc and/or FICZ. (F) FITC-Dextran fluorescence in organoids challenged with 60 µM EDTA for 24 h in combination with 0 or 60 µM zinc with or without 100 nM. for 24 hours. Statistical analysis of the data was performed using 2-way ANOVA followed by either Tukey's multiple comparison tests. Caco-2 cell data are means ± SEM from three independent experiments. n=3 per group. Human ileum organoid data are means ± SEM from one experiment. n=10 per group. *p<0.05, **p<0.01, ***p<0.001, ****p<0.0001, ns not significant.

### Extended Data Figure 4

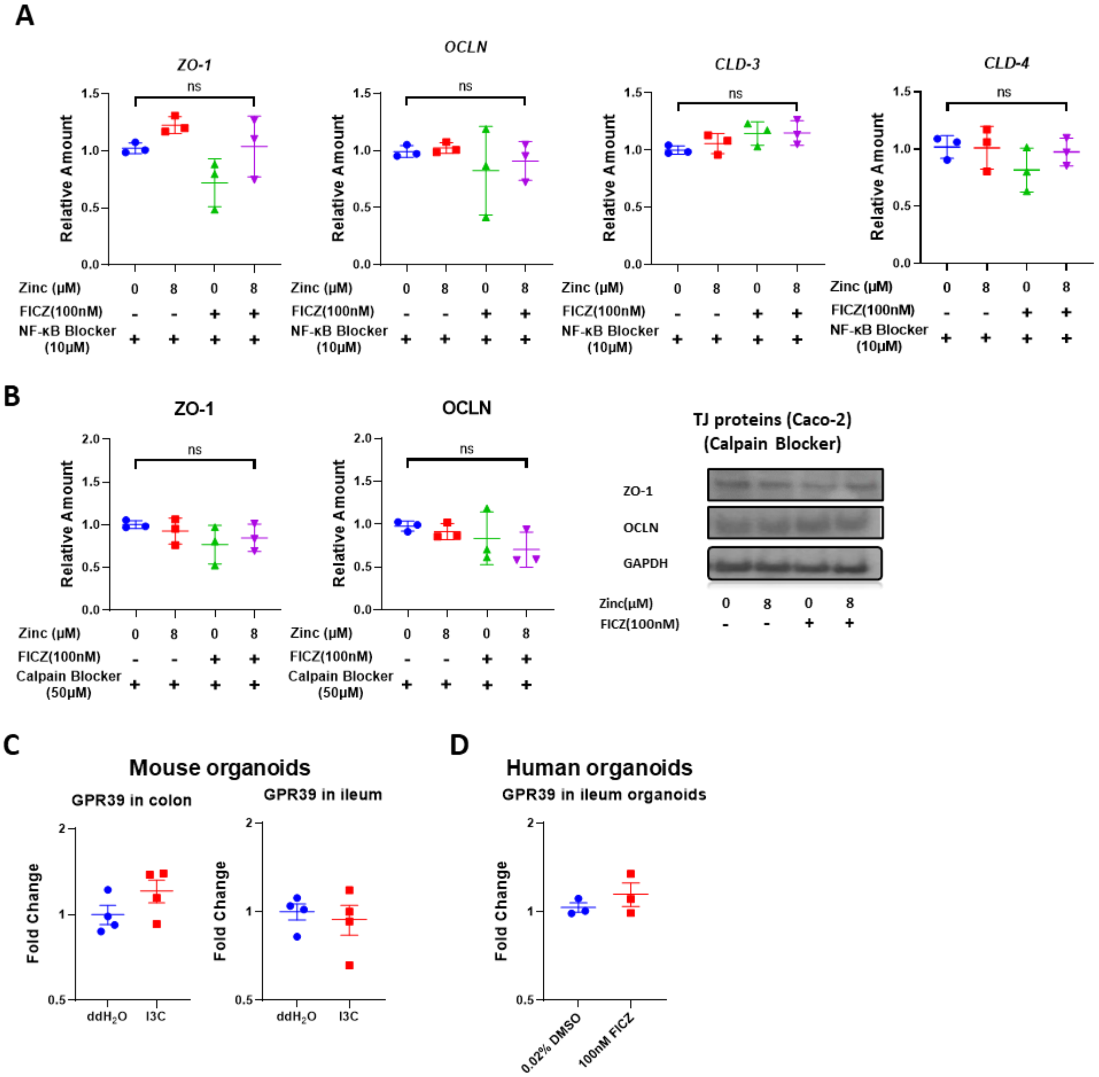


**Extended data Figure 4: Combined treatment of AHR agonist FICZ inhibits activities of NF-κβ and calpains to improve tight junction protein abundance**

The stimulatory effects of zinc and FICZ co-treatment on tight junction proteins and their respective transcripts are lost after blocking of NF-κB or calpain. Abundance of transcripts for TJ proteins in Caco-2 cells treated with zinc and/or FICZ for 24 h after 4 h pre-treatment with (A) 10 µM NF-κB Blocker, QNZ (EVP4593) or (B) 50 µM Calpain Blocker, Calpeptin. (C) Treatment of mouse ileum or colon organoids or (D) human ileum organoids with 100 nM FICZ have no effect on expression of mRNA for the zinc receptor, GPR39. Statistical analysis of the data was performed using unpaired t-tests. Data are means ± SEM from three independent experiments. n=3 per group. *p<0.05, **p<0.01, ***p<0.001, ****p<0.0001, ns not significant.

### Extended Data Figure 5

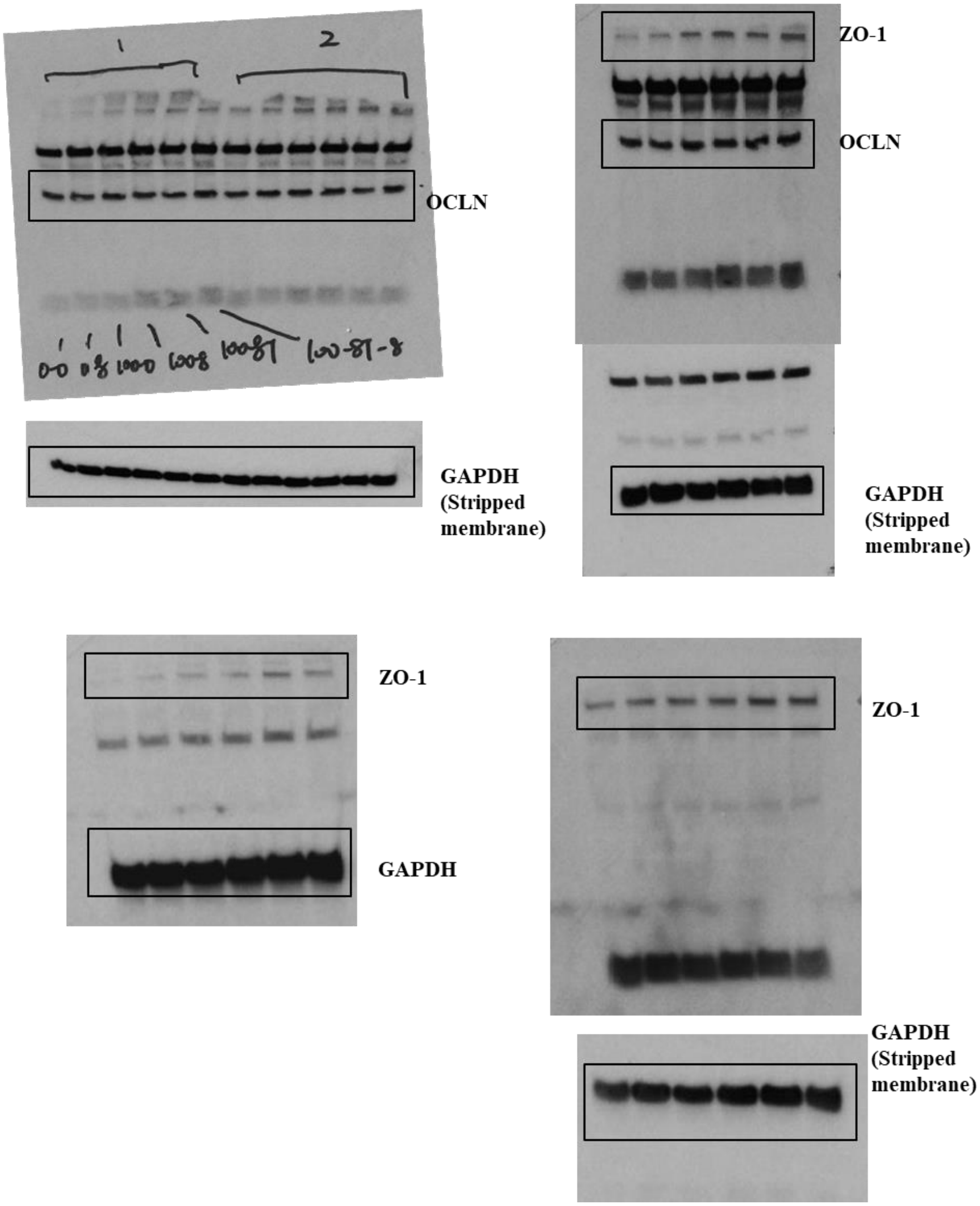


**Extended data Figure 5: Western Blots showing whole blots of ZO-1 and occludin**

Whole Western Blots corresponding to data shown in Figure 3.

### Extended Data Figure 6

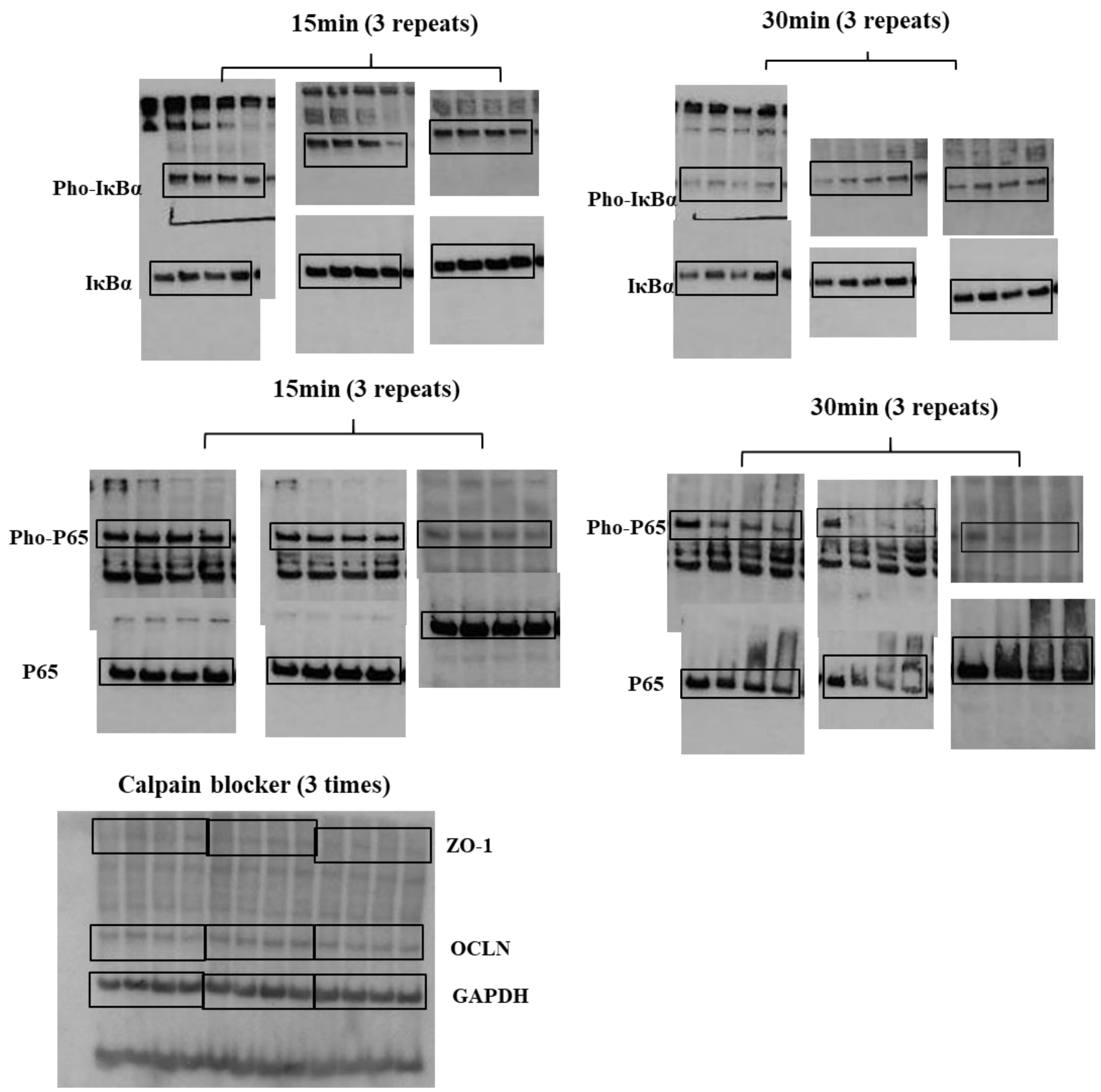


**Extended data Figure 6: Western Blots showing whole blots for phosphorylation of NF-κβ subunits and whole blots for ZO-1 and OCLN**

Whole Western Blots for IκBα$^{ser32/36}$, P65$^{ser536}$, total IκBα, and P65, and ZO-1 and OCLN corresponding to data shown in Figure 4 and 5.

**Extended Data Table 1: qPCR primers used in AHR ChIP-qPCR and location of amplicons and the respective genes.** Genomic locations of genes and PCR fragments are from ENSEMBL human genome assembly GRCh38.p13

| Gene | Alias | Strand | Gene Location | Forward primer | Reverese primer | Fragment location |
|---|---|---|---|---|---|---|
| SLC39A4 | ZIP4 | Reverse | Chr8: 144,409,742-144,416,844 | CCCCAGATCACCGCGGGAGG | CGCGCAGACCTGGTGAGTGG | Chr8: 144,413,632-144,413,761 |
| SLC39A6 | ZIP6 | Reverse | Chr18: 36,108,531-36,129,385 | CGCTTTTCCGTGTGTTCGCCG | TGGCTAGCGAGTAGGGCCTCG | Chr18: 36,129,442-36,129,543 |
| SLC39A7 | ZIP7 | Forward | Chr6: 33,200,305-33,204,439 | CCACCCTCCCCTCAAATCAC | CGTTCCACCTTCGCCCTAAT | Chr6: 33,200,746-33,200,859 |
| SLC39A10 | ZIP10 | Forward | Chr2: 195,575,977-195,737,702 | GTTTGGTACCGAGCGGAGAG | GACTCCCCGGTTCTGTTCTC | Chr2: 195,657,235-195,657,400 |
| MTF1 | MTF-1 | Reverse | Chr1: 37,809,574-37,859,592 | CCTCCAATTGGCGAGCGGGG | CTCCTTGACTCCGGTGCCGC | Chr1: 37,859,711-37,859,839 |

**Extended data Table 2: Primers for qPCR for gene expression analysis.** Probe refers to the identifiers in the Universal Probe Library (UPL).

| *Gene Name* | *Forward Primer* | *Reverse Primer* | *Probe* |
|---|---|---|---|
| *human* | | | |
| *MTF1* | *AAGGGTGTGGTTTCCATCAA* | *GATCCACAAAGACCCCAGTG* | *#44* |
| *MT1A* | *CTTGGGATCTCCAACCTCAC* | *GCATTTGCAGGAGCCAGT* | *#68* |
| *SLC39A2* | *GAACAGATCAGCAAGTGAGAGAAA* | *AGCTCTCCATAGGGATACTCCA* | *#09* |
| *SLC39A4* | *CCTCTTCCTGCTGCACAAC* | *CATCCTCGTACAGGGACAGC* | *#03* |
| *SLC39A6* | *ACTGGCCGTTGGGACTTT* | *ATGGTGGTGACTTGCATGAG* | *#09* |
| *SLC39A7* | *ATGGAGGCTATGGGGAGTCT* | *GGGGATAAGGAAGAGGACAAA* | *#01* |
| *SLC39A10* | *TGAATACACGATTTGGTGCAG* | *TTGTGTGCATATGTACCTTCATTTC* | *#55* |
| *MUC2* | *ACCCACCAGCACACAGAGTA* | *GGGGTTGGGGTTACCGTAT* | *#09* |
| *CLDN1* | *CCTATGACCCCAGTCAATGC* | *ACAGCAAAGTAGGGCACCTC* | *#08* |
| *CLDN3* | *AACCTGCATGGACTGTGAAA* | *GGTCAAGTATTGGCGGTCAC* | *#50* |
| *CLDN4* | *GGGACTGGGCAGAGACTG* | *TTGGGAAGTTGTCCGAGTG* | *#08* |
| *OCLN* | *AGGAACCGAGAGCCAGGT* | *TGAGCAATGCCCTTTAGCTT* | *#84* |
| *ZO-1* | *TGCATGATGATCGTCTGTCC* | *AAGTGTGTCTACTGTCCGTGCTAT* | *#01* |
| *GAPDH* | *AGCCACATCGCTCAGACAC* | *GCCCAATACGACCAAATCC* | *#60* |
| *UBC* | *GGAAGGCATTCCTCCTGAT* | *CCCACCTCTGAGACGGAGTA* | *#11* |
| *Beta-Actin* | *agagctacgagctgcctgac* | *cgtggatgccacaggact* | *#09* |
| *mouse* | | | |
| *Mtf1* | *CCAAGAGACTAGTTGGCAGCA* | *GGTGGGACCAAGATCACCT* | *#10* |
| *Mt1* | *caagtgcacctcctgcaa* | *ttcgtcacatcaggcacag* | *#18* |
| *Slc39A4* | *CAGCTACTGCAGAAGATTGAGG* | *TCCAGCAGTTGGGGAAGAT* | *#07* |
| *Slc39A6* | *CCAGTCCCTTCGGACCTC* | *CTGTGGCCATTGCACCTT* | *#70* |
| *Slc39A7* | *GGATTTTGCCATCCTGGTC* | *TTGCAGTCACGAGTTGCAG* | *#71* |
| *Slc39A10* | *TTTCAGATCATAAGTTAAACAGCACA* | *CCGAGTCATCCGTTCCAG* | *#89* |
| *Ubc* | *GACCAGCAGCAGGCTGATCTT* | *CCTCTGAGGCGAAGGACTAA* | *#11* |

**Extended data Table 3: Antibodies use in Western Blot of cell line and organoids with associated information on dilutions and blocking agent used.**

| *Antibody Name* | Company and Cat No. | Dilution | Blocking agent |
|---|---|---|---|
| *GAPDH* | Sigma-Aldrich, MAB374 | 1:20,000 | 5% skim milk |
| *GAPDH* | Cell Signalling Technology, 2118 | 1:2500 | 5% skim milk |
| *ZO-1* | Cell Signalling Technology, 13663 | 1:800 | 5% skim milk |
| *Occludin* | Santa Cruz Biotechnology, sc-133256 | 1:200 | 5% skim milk |
| *Occludin* | Santa Cruz Biotechnology, sc-133256 | 1:100 | 5 % normal goat serum |
| Occludin | Servicebio, GB111401 | 1:500, | 3% BSA |
| Muc2 | Santa Cruz Biotechnology, sc-515032 AF488 | 1:100 | 5 % normal goat serum |
| Muc2 | Servicebio, GB11344, | 1:500 | 3% BSA |
| *Claudin1* | Invitrogen, 51-9000 | 1:250 | 5% skim milk |
| *Claudin3* | Invitrogen, 34-1700 | 1:250 | 5% skim milk |
| *Claudin4* | Invitrogen,32-9400 | 1:250 | 5% skim milk |
| *Phospho-NF-κB p65* | Cell Signalling Technology, 3033 | 1:1000 | 5% BSA |
| *NF-κB p65* | Cell Signalling Technology, 8242 | 1:1000 | 5% skim milk |
| *Phospho-IκBα* | Cell Signalling Technology, 9246 | 1:1000 | 5% BSA |
| *IκBα* | Cell Signalling Technology, 4814 | 1:1000 | 5% skim milk |
| *Anti-Mouse IgG* | Bio-rad, 1705047 | *1:10,000* | 5% skim milk |
| *Anti-Rabbit IgG* | Santa Cruz Biotechnology, sc-2357 | *1:4000* | 5% skim milk |
| Goat anti-Mouse IgG | Invitrogen, A-11031 | 1: 250 | 5 % normal goat serum |
| Goat Anti-rabbit IgG | Servicebio, GB1213 | 1: 200 | 3% BSA |
